# A signaling-optimized TIGIT chimeric switch receptor promotes metabolic fitness and antitumor efficacy of TCR/CAR T cells

**DOI:** 10.64898/2026.10.02.755760

**Authors:** Edite Antunes Ferreira, Kevin Jan Legscha, Ulrike Regina Schmits, Alexandra Reinbrecht, Matthias Theobald, Hakim Echchannaoui

## Abstract

Adoptive T cell therapies have revolutionized the treatment of hematological malignancies, however, their efficacy against solid tumors often remains limited. Lack of activation and overabundance of immune checkpoint-mediated inhibitory signals from cancer cells drive T cell dysfunction and limit persistence. To circumvent this imbalance towards inhibitory factors, we engineered a chimeric switch receptor (CSR) that repurposes the immunosuppressive TIGIT extracellular domain by fusing it to a CD28-derived stimulatory signaling tail. Here we show that while the prototype CSR design enhances immediate cytolytic activity, it also induces premature proliferative arrest and terminal differentiation due to excessive tonic signaling. By introducing a strategic point mutation within the CD28 cytoplasmic domain, we “fine-tuned” the activation threshold of the CSR for optimal functional properties. Engineered T cells with “fine-tuned” CSR exhibit superior mitochondrial bioenergetics and sustained metabolic flux in an *in vitro* system of chronic antigen exposure. In preclinical xenograft models, fine-tuned CSR-expressing T cells demonstrated robust expansion and durable persistence, leading to complete tumor eradication. These findings provide a scalable strategy to rewire inhibitory checkpoints into tailored costimulatory signals, offering a potent approach to enhance the metabolic fitness and persistence of therapeutic T cells, in suppressive tumor microenvironments, in particular.

**Graphical Abstract:** 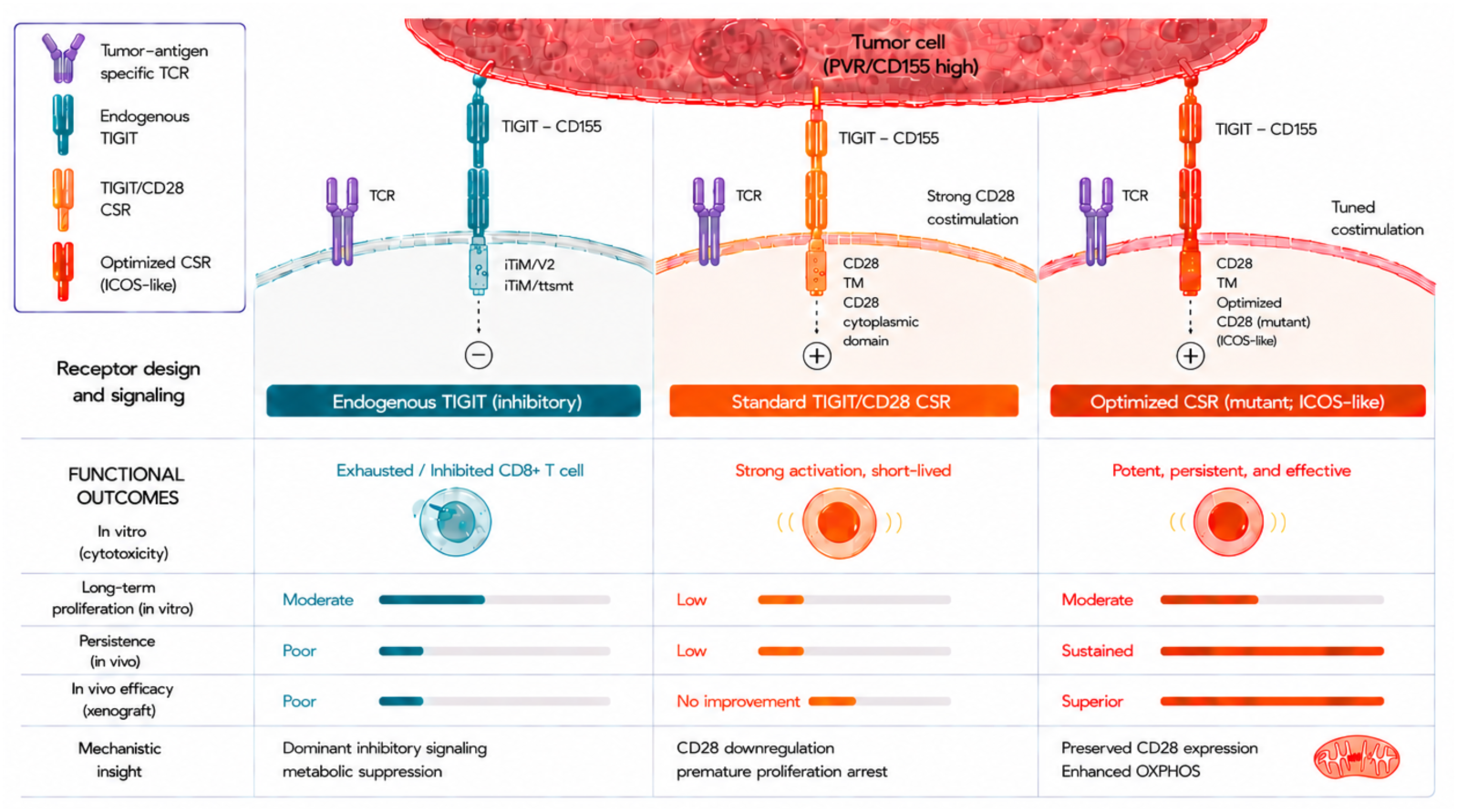

**Significance:** This work shows that inhibitory signals as transmitted by tumor cells can be converted into controlled activating signals to improve T cell cancer therapies. By fine-tuning the engineered CSR, we demonstrated longer-lasting anti-tumor activity, without driving T cell exhaustion, offering a promising alternative to standard TIGIT checkpoint inhibitors for cellular therapy of solid cancer.

## Introduction

Adoptive transfer of T cells genetically engineered with a tumor-antigen specific T cell receptor (TCR), or chimeric antigen receptor (CAR) has revolutionized cancer immunotherapy and established itself as one of the most effective approaches to treat otherwise refractory hematological malignancies, however, it has achieved limited success against solid tumors ^1–3^. A key factor limiting the efficacy of cellular therapy in clinical use for solid cancers is the dysfunction and insufficient expansion of infused T cells, driven by a predominance of inhibitory over co-stimulatory signals within the tumor microenvironment ^4,5^. Therefore, strategies ensuring sufficient co-stimulation while avoiding inhibitory signals are essential for successful TCR- and CAR-based T cell therapy against solid tumors.

Inhibitory receptors, such as programmed cell death protein 1 (PD-1), cytotoxic T-lymphocyte-associated Protein 4 (CTLA-4), and T cell immunoreceptor with Ig and ITIM domains (TIGIT), were identified as powerful immune checkpoints that impede anti-tumor responses of adoptive cell therapy (ACT) ^5–7^. Different strategies interfering with these pathways were developed to increase T cell activation and anti-tumor immunity. Blockade of inhibitory receptors with monoclonal antibodies (immune checkpoint inhibition = ICI) was among the first immunotherapeutic approaches to demonstrate clinical benefits and is now widely used ^8^. An alternative option to circumvent inhibitory signaling and switch it to co-stimulation is the use of so-called chimeric switch receptors (CSR). CSRs are composed of the extracellular domain of an inhibitory and the intracellular domain of a co-stimulatory receptor. Binding of CSRs to their ligands provides activating instead of inhibitory signals and, hence could improve proliferation, persistence and cytotoxicity of CAR- and TCR-engineered T cells ^9^.

First designed CSRs were based on the extracellular domain of PD-1 and the co-stimulatory intracellular domain of CD28 and reported substantial response rates and durable remissions in CAR T cell therapies in different B cell-derived malignancies ^10–12^. Expression of PD-1/CD28-based CSRs resulted in enhanced anti-tumor activity of TCR-^13^, as well as of CAR-engineered T cells ^14^. First clinical phase 1 trials with PD1/CD28-CAR T cells have already been initiated and have reported objective responses even in B-cell lymphoma patients that experienced relapse after conventional CD19-CAR T cell therapy ^15^, highlighting the therapeutic potential of CSRs.

While most monoclonal antibodies and CSRs have been designed to target PD-1 or CTLA-4, relatively few studies have reported utilizing extracellular domains of other inhibitory receptors for CSR engineering, such as T-cell immunoglobulin mucin domain molecule 3 (TIM-3) or TIGIT ^16,17^. TIGIT has gained significant attention as a target for immunotherapy due to its inhibitory effects on T cell and NK cell function ^18,19^, as well as its regulatory role in other immune cells, including B cells ^20,21^. Upon activation, TIGIT suppresses T cell proliferation, diminishes cytokine production, and compromises anti-tumor immunity ^22,23^. Its main ligands, CD155 and CD112, are widely expressed on tumor cells ^24,25^, while high TIGIT expression is observed on T cells in cancer patients ^26,27^. Furthermore, it was shown in preclinical studies, that blockade of TIGIT by monoclonal antibodies could restore anti-tumor immunity of T cells ^27^.

These characteristics turn TIGIT into an attractive candidate for CSR design, potentially enabling redirection of its immunosuppressive properties into activation signals for engineered T cells. Arming T cells with a TIGIT CSR could allow selective engagement with tumor-associated ligands and reprogramming of T cells to resist the immunosuppressive tumor microenvironment-mediated T cell exhaustion. This strategy may provide a novel approach for improving adoptive cell therapy outcome by leveraging TIGIT’s tumor-specific interactions to boost therapeutic efficacy.

In this study, we engineered a TIGIT-based CSR comprising the extracellular domain of TIGIT fused to the transmembrane and cytoplasmic domains of CD28 (TIGIT/CD28, thereafter TIGIT/28). While the expression of TIGIT/28 CSR enhanced the anti-tumor cytotoxicity of TCR-engineered T cells *in vitro*, *in vivo* therapeutic efficacy was moderate. To address this limitation, we introduced a single amino acid substitution in the YMNM motif of the CD28 intracellular domain, replacing it with the YMFM motif present in the ICOS co-stimulatory domain. This modification effectively mitigated CD28-associated T cell exhaustion and demonstrated improved effector function, *in vivo* persistence, and anti-tumor immunity in both, TCR- and CAR-engineered T cells.

Our findings highlight the importance of fine-tuning the intracellular signaling domain of a CSR for optimal T cell effector functions, offering a promising strategy for ACT outcome.

## Results

### Stable expression of TIGIT/28 CSR on TCR T cells

The inhibitory immune checkpoint TIGIT is upregulated upon T cell activation and strongly expressed by (highly) differentiated and exhausted T cells ^23,28^. Its ligand, CD155, is expressed by tumor cells in a wide range of human cancers ^24,26^. TIGIT can significantly inhibit T cell cytotoxicity ^7^ and its blockade via monoclonal antibodies was shown to improve T cell immunity against cancer in vitro and in vivo ^27^. We constructed a TIGIT-based chimeric switch receptor to not only prevent T cell inhibition, but to use the CD155/TIGIT interaction to provide a co-stimulatory signal mediated by the intracellular domain of CD28 to further enhance T cell function. Therefore, the signal peptide (amino acid 1-21) and extracellular domain (amino acid 22-141) of TIGIT were fused to the transmembrane (amino acid 153-179) and cytoplasmic domain (amino acid 180-220) of CD28 (**Fig. 1A**).

**Figure 1:**
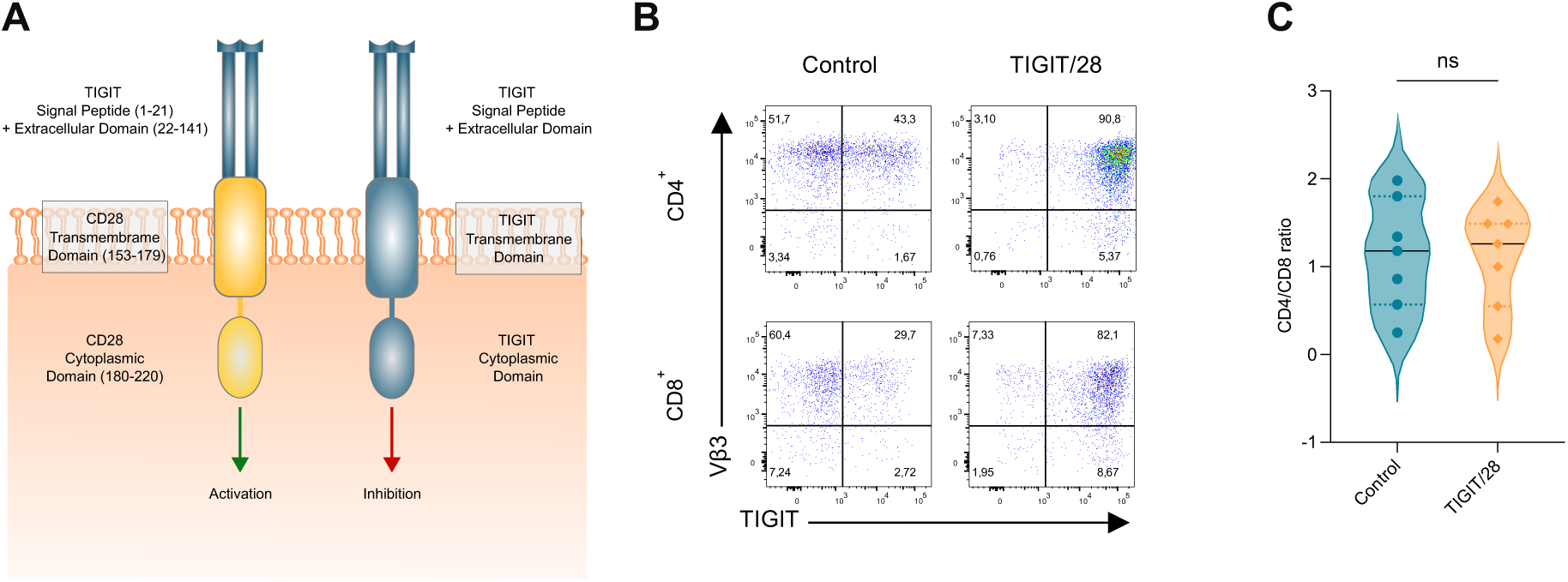
Design and expression of the TIGIT/28 chimeric switch receptor on TCR T cells. **(A)** Graphical illustration of the chimeric switch receptor design and the regular TIGIT protein including the extracellular, transmembrane and cytoplasmic domain. **(B)** Representative pseudocolor plots showing expression levels of TIGIT and the co-transduced scTCR (Vβ3) in TIGIT/28 + TCR and TCR + empty vector (control) T cells. **(C)** Flow cytometry analysis depicted as violin plot showing the CD4/CD8 ratio of TCR-transduced T cells without additional CSR (control) and with CSR (TIGIT/28) (n=7 donors). Median is shown. To test statistical significance against the control in C, the *p*-value was calculated using the two-tailed Student’s t-test (*p*=0.6573, t=0.4665, degree of freedom=6). n.s. = not significant; * *p* < 0.05; ** *p* < 0.01; *** *p* < 0.001.

To test the function and impact of this TIGIT/28 CSR on human T cells in the context of T cell-based cancer immunotherapy, primary T cells from healthy donors were genetically modified with a tumor antigen-specific single-chain TCR (scTCR) and the CSR (hereafter TIGIT/28 TCR^+^ T cells). Control cells were modified with the scTCR and an empty vector (referred to as ‘control TCR T cells’, if not otherwise specified) ^29^. Expression of both receptors, CSR and TCR, was determined by flow cytometry. The CSR was stained using an anti-TIGIT antibody. The recombinant TCR was stained using a murine Vβ3 antibody. After selection, TCR expression was >90% in both TIGIT/28 TCR^+^ T cells and control TCR T cells (Vβ3 expression > 90%, **Fig. 1B**). The CSR was detected in over 90% of TIGIT/28 TCR^+^ T cells, while control TCR T cells showed varying endogenous expression of TIGIT (**Fig. 1B**). Overexpression of the CSR did not alter the ratio between CD4^+^ and CD8^+^ T cells compared to control cells (**Fig. 1C**). Taken together, our designed TIGIT/28 CSR is stably co-expressed together with a scTCR on healthy donor T cells and does not modify expression of the transgenic TCR and the CD4/CD8 ratio as compared to control TCR T cells.

### Co-expression of TIGIT/28 CSR improves anti-tumor toxicity of TCR T cells and correlates with enhanced cytokine response

Effector functions of TIGIT/28 TCR^+^ T cells were compared to control TCR T cells by measuring tumor-specific lysis, degranulation capacity and IFN-γ secretion. The osteosarcoma cell line Saos2/143, which expresses the high-binding capacity TIGIT ligand CD155 and the p53_264-272_ antigen recognized by the utilized scTCR construct in this study, served as a target model for functional assays ^23,30^. In a short-term chromium-release assay, T cells and target cells were co-cultured at different effector to target ratios for 5 h. Here, control and TIGIT/28 TCR^+^ T cells showed a comparable specific tumor lysis (**Suppl.-Fig.1**). We previously reported that prolonged co-culture (24h) of antigen-specific T cells with target tumor cell line Saos2/143 induced a marked increase in TIGIT expression, which was associated with impaired tumor cell lysis ^30^. To assess the long-term cytotoxicity of TIGIT/28 TCR^+^ T cells, we performed tumor colony-forming assays with three consecutive rounds of 24 h co-culture. In this repetitive co-culture experiment, tumor lysis by control TCR T cells declined, resulting in a high tumor cell viability (mean 59.2%). In contrast, TIGIT/28 TCR^+^ T cells demonstrated an efficient anti-tumor response resulting in a low or moderate tumor cell viability (mean 21.9%, **Fig. 2A**).

**Figure 2:**
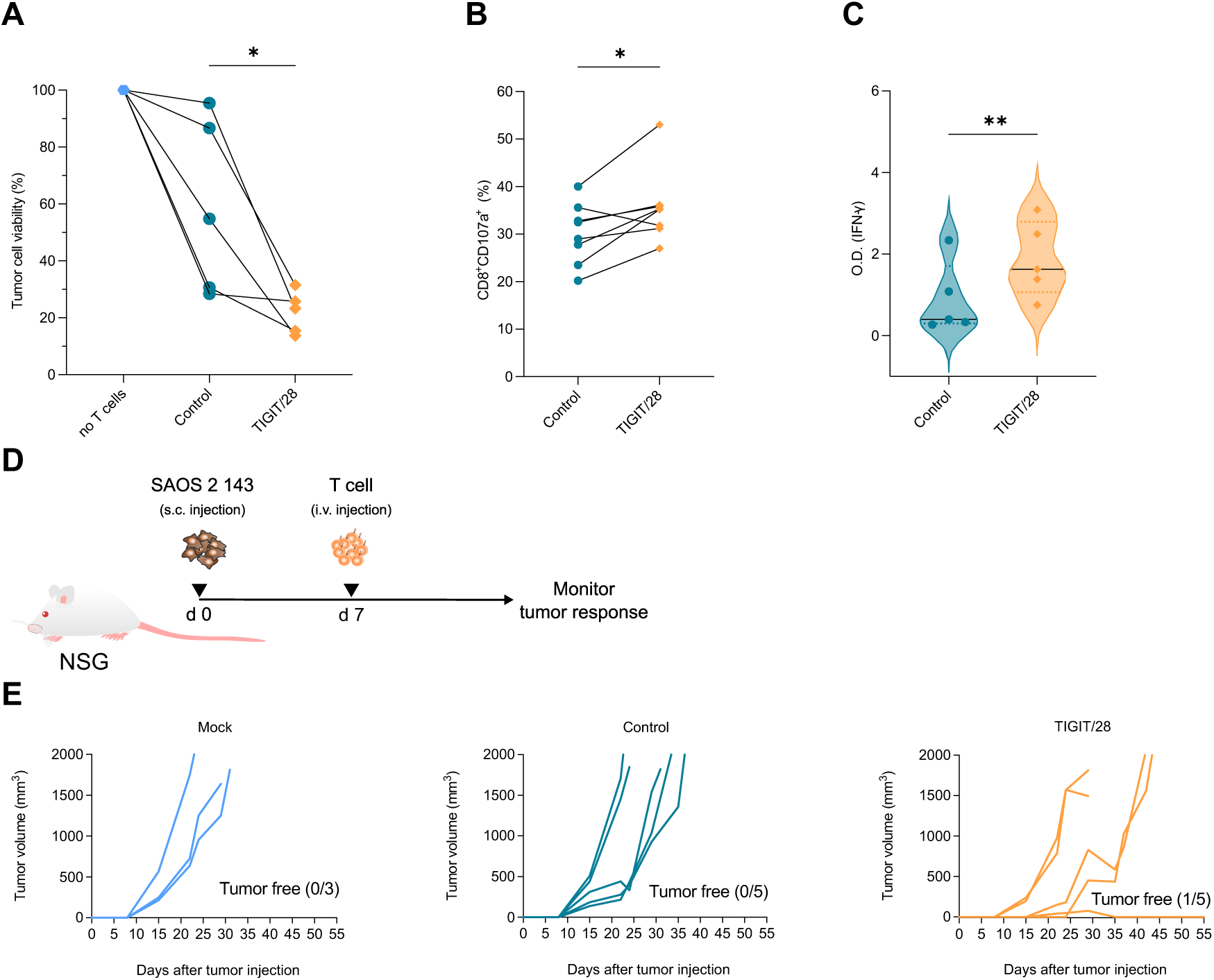
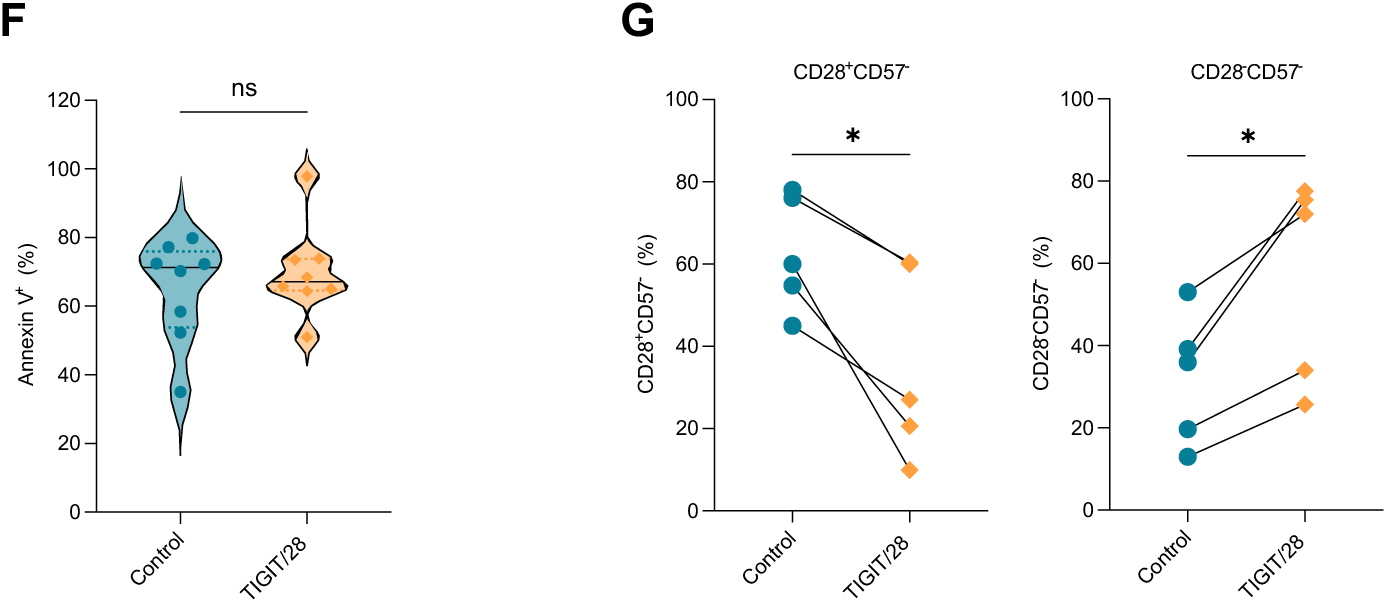
Characterization of TIGIT/28 CSR functions *in vitro* and in xenograft tumor model. (**A** - **C**) T cells were engineered to express tumor-antigen specific scTCR +/-CSR (TIGIT/28 vs. empty vector control) and cultured with or without TCR-specific tumor cells for three consecutive rounds of 24 h each (total 96 h). **(A)** Tumor cell viability in percentage (*p*=0.0428, n=5 biological replicates). **(B)** Degranulation capacity of CD8^+^ T cells indicated by expression of CD107a, measured by flow cytometry (*p*=0.0244, n=8 biological replicates). **(C)** Violin plots demonstrating secretion of IFN-γ, reported by optical density using enzyme-linked immunosorbent assay (ELISA) (*p*=0.0053, n=5 biological replicates). **(D)** Experimental design of the xenograft model to evaluate anti-tumor responses *in vivo*. **(E)** Tumor (Saos2/143) growth over time after subcutaneous injection on day 0 and intravenous T cell transfer on day 7 (n=3-5 animals per group: unmodified (mock) T cells, TCR + empty vector T cells (control) and TCR + TIGIT/28 T cells. **(F)** Apoptosis rate upon stimulation, indicated by positivity for Annexin V in flow cytometry (*p*=0.4853, n=8 biological replicates). **(G)** Frequency of CD28^+^CD57^-^ (left) and of CD28^-^CD57^-^ (right) T cells, measured by flow cytometry (*p*=0.0147 and *p*=0.0135, n=5 biological replicates). To test statistical significance against the control in A-C and F-G, *p*-values were calculated using the two-tailed Student’s t-test. n.s. = not significant; * *p* < 0.05; ** *p* < 0.01; *** *p* < 0.001.

We examined the expression of the lysosomal-associated protein CD107a (=LAMP-1) upon tumor stimulation (one round of co-culture) as an indicator of the capacity of T cells to secrete toxic granules such as granzymes and perforin ^31^. Further, we determined IFN-γ secretion by ELISA. After co-culture with Saos2/143 tumor cells (three rounds), analysis of CD8^+^ control TCR T cells displayed a mean expression of 35.1%, while 40.0% of TIGIT/28 TCR^+^ T cells showed expression of CD107a (**Fig. 2B**). Additionally, secretion of IFN-γ was markedly enhanced in TIGIT/28 TCR^+^ T cells compared to control TCR T cells (mean O.D. control 0.89 vs. TIGIT/28 1.89, **Fig. 2C**). Taken together, the enhanced tumor killing capacity of TIGIT/28 TCR^+^ T cells correlates with increased expression of CD107a and IFN-γ secretion *in vitro*.

### TIGIT/28 TCR T cells exhibit limited activity in xenograft tumor model

To explore the functionality of TIGIT/28 TCR^+^ T cells, we used a xenograft *in vivo* tumor model in which NSG mice were engrafted with the osteosarcoma tumor cell line Saos2/143. Seven days after subcutaneous injection of tumor cells, T cells engineered with either scTCR and TIGIT/28 CSR (TIGIT/28 TCR^+^ T cells) or scTCR and empty vector (control TCR T cells) were injected intravenously. Mice injected with unspecific (mock) T cells were used as an additional control. Mice received a similar T cell composition, with regard to CD4/CD8 ratio and TCR expression (**Suppl.-Fig. 2A**). Tumor growth was monitored over time (**Fig. 2D**). Control TCR T cells induced a minor delay in tumor growth compared to unspecific T cells but failed to eliminate the tumor. TIGIT/28 TCR^+^ T cells further delayed the tumor growth but did not result in complete tumor eradication (except in one out of five mice) (**Fig. 2E**). The overall therapeutic effect of TIGIT/28 CSR *in vivo* was limited with four out of five mice developing growing tumors. Human CD3^+^ T cells were detected in all animals 14 days after T cell injection, with no significant difference between the treated groups (**Suppl.-Fig. 2B**). However, on day 27 after injection, no T cells were detected in the peripheral blood except for the mouse, which showed complete tumor control, yet presence of circulating T cells in this animal was just transient as no T cells could be detected at later time points. To better understand the poor *in vivo* persistence and thus anti-tumor response of CSR-T cells, we analyzed additional features, including apoptosis rate and differentiation status of these T cells after three to four rounds of *in vitro* stimulation. The apoptosis rate was similar in both control and TIGIT/28 CSR-T cells, as assessed by the frequency of Annexin V^+^ cells (**Fig. 2F**). Analysis of CD28 and CD57 expression (as critical markers of T cell differentiation) showed an increase in early differentiated CD28^-^CD57^-^ T cells and reduced frequency of naïve CD28^+^CD57^-^ population in TIGIT/28 TCR T cells (**Fig. 2G**) suggesting a premature differentiation process that might contribute to the poor persistence and insufficient anti-tumor responses *in vivo*.

### Optimized TIGIT/28 CSR enhances T cell anti-tumor activity *in vitro* while maintaining proliferative capacity and preventing premature differentiation

Studies in CAR T cells already demonstrated that CD28 co-stimulation can promote T cell exhaustion and therefore contributes to short persistence *in vivo* ^32^. Guedan et al. have shown that a single amino acid residue (asparagine (N) in the YMNM motif) of the CD28-costimulatory domain facilitates CAR T cell exhaustion ^33^. In this study, changing this single amino acid asparagine (N) to phenylalanine (F) to create a YMFM motif found in the Inducible T-cell Costimulator (ICOS)-costimulatory domain, resulted in increased persistence and improved anti-tumor immunity. Therefore, we aimed to apply this optimization to our chimeric switch receptor construct and test its effect on T cell proliferation and functionality *in vitro* and *in vivo*. Using site-directed mutagenesis, an amino acid change from asparagine to phenylalanine at position 182 was induced to turn the YMNM motif of CD28 to the YMFM motif of ICOS (**Suppl.-Fig. 3A**). The successful amino acid change was confirmed by sequencing of the CSR construct. T cells engineered with the N182F mutated TIGIT/28 CSR (thereafter TIGIT/28F) showed comparable expression of TIGIT to T cells engineered with the TIGIT/28 (mean expression 89.7% vs 92.9%, **Fig. 3A**) and higher expression compared to the endogenous TIGIT (Control CD8^+^: mean expression 19.7%). The ability of TIGIT/28F TCR T cells to expand upon stimulation was tested *in vitro*. T cells engineered with the scTCR and the TIGIT/28F CSR exhibited a slightly improved proliferative capacity compared to T cells expressing the scTCR and the TIGIT/28 CSR (mean PDL: 7.5 vs. 6.0, **Fig. 3B**). TIGIT/28 TCR T cells also had a statistically significant lower PDL as compared to control TCR T cells (mean PDL: 6.0 vs. 8.1), while no significant difference was observed between TIGIT/28F and control TCR T cells (mean PDL: 7.5 vs. 8.1, **Fig. 3B**). This preserved proliferation was further associated with an unaltered cellular phenotype of TIGIT/28F CSR compared to control TCR T cells, while co-expression of the TIGIT/28 CSR again promoted differentiation from CD28^+^CD57^-^ (mean: 34.9%) towards CD28^-^CD57^-^ T cells (mean: 61.4%), TIGIT/28F CSR showed similar levels of differentiation to the control values (**Fig. 3C**).

**Figure 3:**
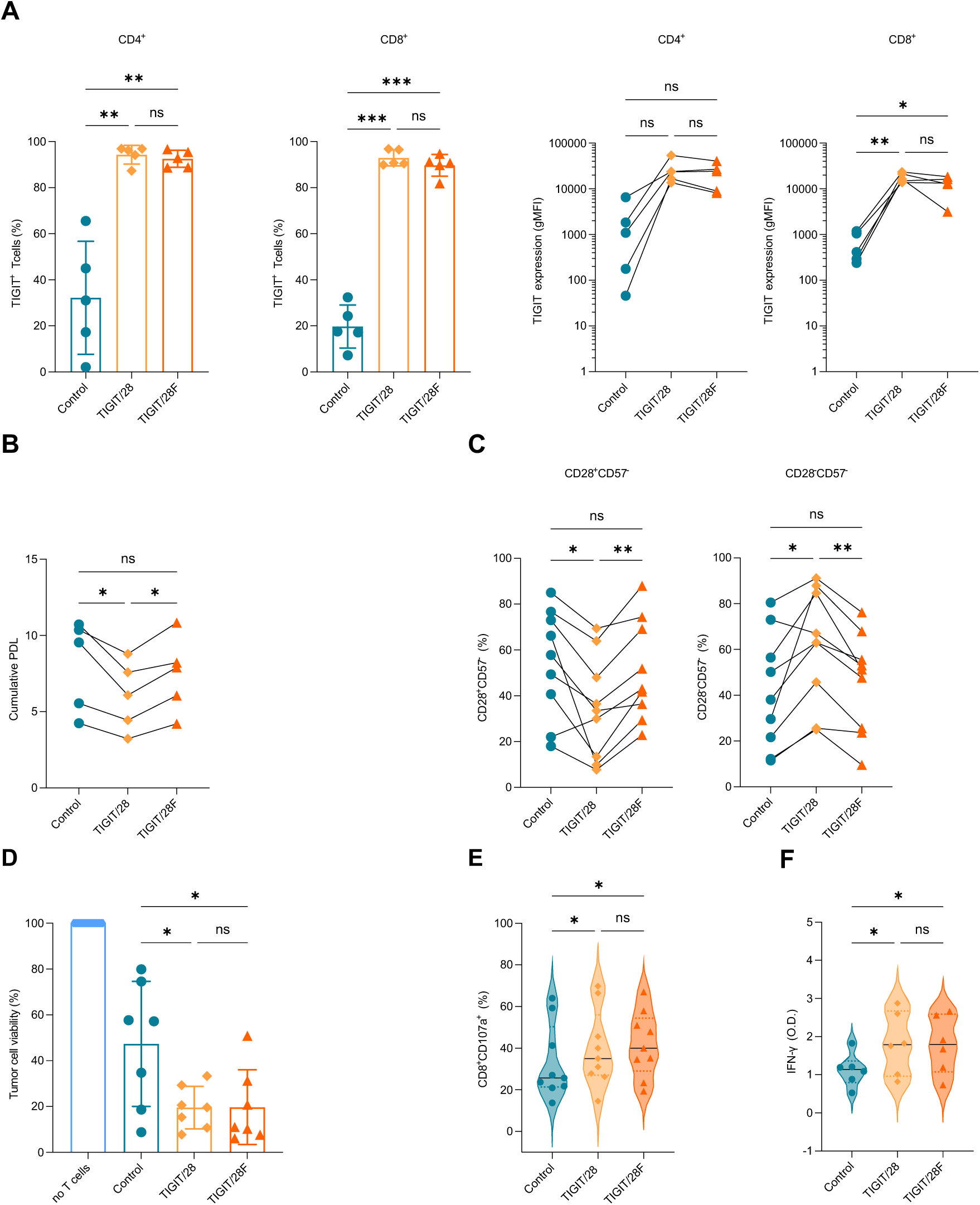
Functional characteristics of the optimized TIGIT/28F CSR *in vitro*. **(A)** Flow cytometry plots showing the percentage of TIGIT positive T cells (left) and the expression intensity of TIGIT per cells in MFI (right) in TCR only (control), TIGIT/28 and TIGIT/28F CD4^+^ and CD8^+^ T cells (CD4^+^ (left): *p*=0.0089, *p*=0.0067 and *p*=0.5884, CD8^+^ (left): *p*=0.0002, *p*=0.0003 and *p*=0.5535, CD4^+^ (right): *p*=0.0554, *p*=0.0513 and *p*=0.3584, CD8^+^ (right): *p*=0.0015, *p*=0.0182 and *p*=0.2409, each n=5 biological replicates). **(B)** Cumulative PDL after *in vitro* culture of TCR T cells +/- CSR (*p*=0.0352, *p*=0.5897 and *p*=0.0127, n=5 biological replicates). **(C)** Frequency of CD28^+^CD57^-^ (left, *p*=0.0232, *p*=0.8177 and *p*=0.0011) and of CD28^-^CD57^-^ (right, *p*=0.0225, *p*=0.7464 and *p*=0.0016) T cells, measured by flow cytometry (each n=9 biological replicates). **(D)** Viability of tumor cell (in percentage) as determined by tumor colonies forming assay after three consecutive rounds of T cell co-culture. Values were normalized to control wells containing the tumors without T cells (*p*=0.0363, *p*=0.0278 and *p*=0.9993, n=6 biological replicates). **(E)** T cell degranulation indicated by expression of CD107a after co-culture with target tumor cells Saos2/143 (*p*=0.0142, *p*=0.0496 and *p*=0.6025, n=9 biological replicates). **(F)** Violin plots representing secretion of IFN-γ, measured by ELISA (*p*=0.0365, *p*=0.0254 and *p*=0.65739185, n=6 biological replicates). Statistical significance in A-F was determined by one-way ANOVA and Tukey’s multiple comparisons test. n.s. = not significant; * *p* < 0.05; ** *p* < 0.01; *** *p* < 0.001.

Next, long-term cytotoxicity assays were performed to evaluate the influence of the modified co-stimulatory domain of the CSR on TCR T-cell-mediated anti-tumor responses *in vitro*. T cells were co-cultured with Saos2/143 for 24h across three consecutive cycles to monitor sustained functional activity. Co-expression of TIGIT/28F CSR resulted in an improved cytotoxicity (mean tumor cell viability 19.7%) compared to control TCR T cells (mean 47.4%). No significant difference was observed between TIGIT/28F and TIGIT/28 CSRs (mean 19.7% vs. 19.5%, **Fig. 3D**). Both TIGIT/28F and TIGIT/28 TCR T cells, showed modestly enhanced CD107a expression (mean 39.6% and 41.8%) compared to the control TCR T cells (mean 33.1%, **Fig. 3E**). Accordingly, secretion of IFN-γ by TIGIT/28F TCR T cells (mean O.D: 1.79) and TIGIT/28 TCR T cells (mean O.D. 1.82) was enhanced (**Fig. 3F**). In short-term killing assays, the EC50 of the scTCR was measured at different peptide concentrations. T cells expressing the TIGIT/28F and TIGIT/28 TCR T cells showed comparable EC50 values compared to control TCR T cells (**Suppl.-Fig. 3B**), demonstrating that the TCR functional avidity remains unaltered in short-term anti-tumor responses, when co-stimulation has less impact. To further confirm the impact of TIGIT and the CSR signaling on the anti-tumor responses, we performed a TIGIT blocking assay. TCR T cells co-expressing TIGIT/28F or TIGIT/28 CSRs were co-cultured with CD155 expressing target tumor cells Saos2/143 in the absence or presence of a monoclonal antibody blocking TIGIT. Blockade of TIGIT resulted in enhanced tumor lysis mediated by control TCR T cells, as they only express the inhibitory endogenous TIGIT receptor. In the first round of co-culture, TIGIT blockade did not impact the cytotoxicity of CSR expressing TCR T cells (**Suppl. -Fig. 3C**). However, after three consecutive rounds of co-culture, TIGIT/28F or TIGIT/28 TCR T cells exhibited an increased anti-tumor response compared to control TCR T cells, which was abrogated in the presence of TIGIT blockade (**Suppl. -Fig. 3C**). As an additional control, we engineered T cells with the CSRs only (without scTCR) to exclude any potential unspecific CSR-mediated lysis of tumor cells (**Suppl.-Fig. 3D**).

Taken together, these findings indicate that modifying the co-stimulatory domain of the CSR prevents premature proliferation arrest while maintaining a less-differentiated T-cell phenotype. This modification results in superior anti-tumor response *in vitro*. Additionally, TIGIT blockade underscores the direct contribution of the CSR to the observed anti-tumor activity.

### TIGIT/28F CSR empowers anti-tumor response and persistence of TCR T cells in xenograft tumor model

To further validate the effective cytotoxicity and sustained proliferative capacity of TIGIT/28F T cells observed *in vitro*, their anti-tumor activity was assessed *in vivo* using the previously described osteosarcoma xenograft model. NSG mice were injected (subcutaneous) with Saos2/143 tumor cells at day 0, followed by T cell injection (intravenous) along with IL-2 (intraperitoneal) (**Fig. 4A**). Groups included animals receiving unspecific (mock), TCR + empty vector (control), TCR + TIGIT/28 or TCR + TIGIT/28F T cells. Mice received similar T cells composition, with regard to CD4/CD8 ratio and TCR expression (**Suppl.-Fig. 4A**). Expression of the TIGIT CSRs was also verified prior ACT transfer (**Suppl.-Fig. 4A**). Mice receiving unmodified mock T cells showed a rapid tumor growth (0/5 tumor free animals, **Fig. 4B**). Control T cells also failed to induce sufficient tumor control (0/5 tumor free animals, **Fig. 4B**). Co-expression of TCR and TIGIT/28 CSR resulted in limited anti-tumor responses, achieving complete tumor eradication in only one out of five mice (1/5 tumor free, **Fig. 4B**). These results are consistent with the outcomes observed in the initial *in vivo* experiment (**Fig. 2E**). Notably, tumors treated with TIGIT/28F TCR T cells were eradicated after initial growth in all mice and remained undetectable throughout the observation period (5/5 tumor free, **Fig. 4B**). After complete tumor eradication some mice treated with TIGIT/28F T cells exhibited signs of tumor-unrelated illness, including body weight reduction, hair loss and apathy, possibly as a consequence of T cell persistence/activation (>50 days after tumor injection). Nevertheless, the anti-tumor response of TIGIT/28F TCR T cells showed a profound survival benefit of treated mice. The median survival of mice treated with TIGIT/28F TCR T cells was increased (64 days) compared to mock (34 days), control (29 days) and TIGIT/28 (32 days, **Fig. 4C**) groups.

**Figure 4:**
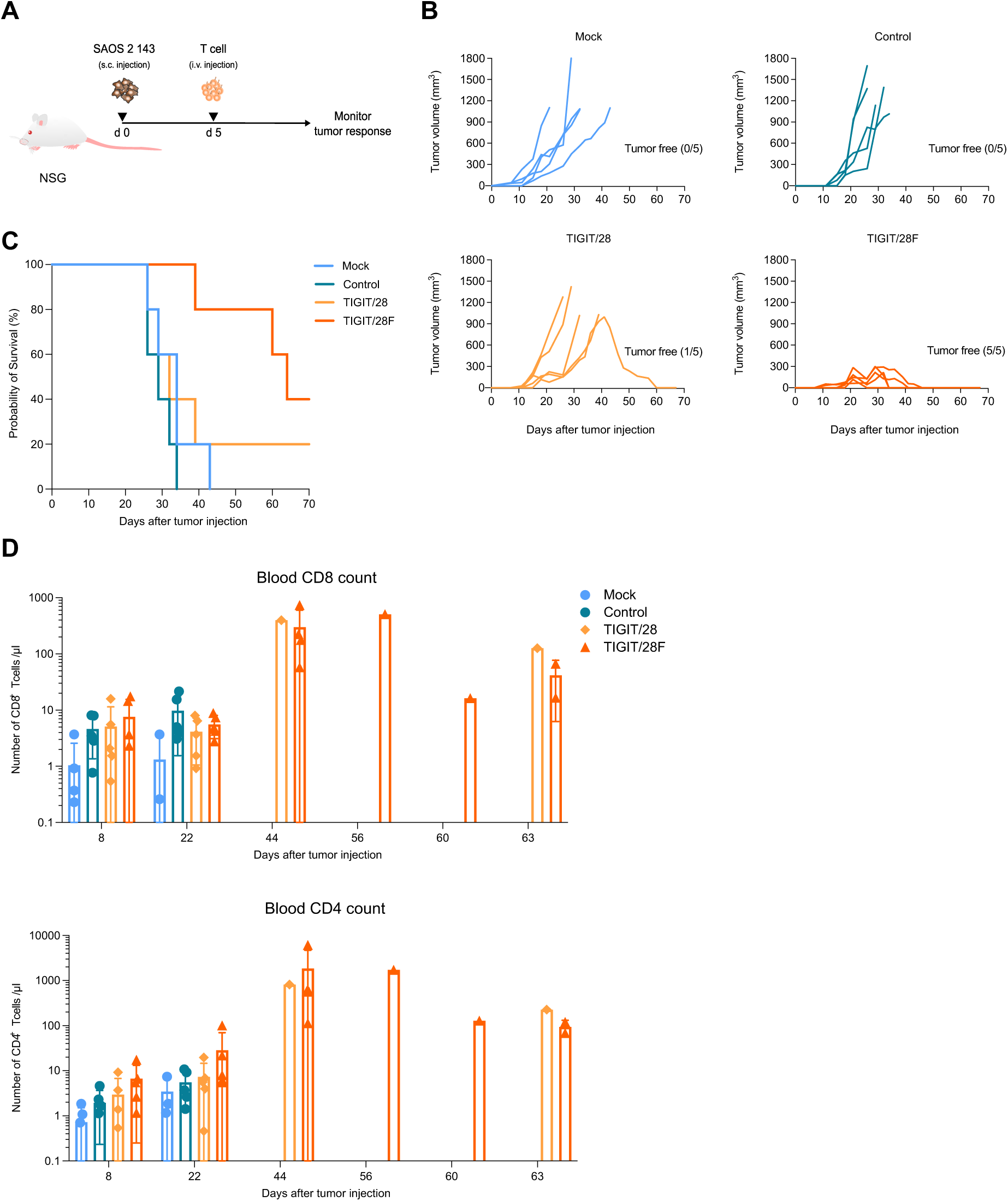
Evaluation of the anti-tumor function of the optimized TIGIT/28F CSR *in vivo*. **(A)** Experimental design of the xenograft model to evaluate anti-tumor responses *in vivo*. **(B)** Tumor volume from the day of tumor injection (day 0) until day 70 (n=5 animals per group: unmodified (mock) T cells, TCR + empty vector T cells (control), TCR + TIGIT/28 T cells and TCR + TIGIT/28F T cells. **(C)** Kaplan-Meier curves demonstrating survival of mice after tumor injection in percentage. **(D)** Counts of human CD8^+^ and CD4^+^ T cells per µl blood as measured by flow cytometry at indicated time points. Statistical significance in C was determined by Log-rank (Mantel-Cox) test (*p*=0.0177). Next, groups were tested for statistical significance against the control with Holm-Šídák’s multiple comparisons test (Control vs. Mock *p=0.3761*, Control vs. TIGIT/28 *p*=0.3761, Control vs. TIGIT/28F *p*=0.00351). n.s. = not significant; * *p* < 0.05; ** *p* < 0.01; *** *p* < 0.001.

As T cell persistence *in vivo* is often correlated with efficient and sustained anti-tumor responses ^42,43^, T cell numbers in the peripheral blood were monitored over time. Infused human CD8^+^ T cells were detected as early as day 8 after injection in nearly all treated mice (**Fig. 4D**). Mock T cells exhibited poor persistence, being detected in only two out of five mice by day 22, while control and CSR-transduced T cells maintained detectable levels in all mice up to this point. By day 44, TIGIT/28 and TIGIT/28F treated mice with tumor eradication showed high and persistent levels of circulating CD8^+^ T cells. Remarkably, TIGIT/28F CD8^+^ T cells expanded in four out of five mice between days 22 and 44, with cell counts ranging from 57.9 - 737.7/µl blood. The expansion and persistence of T cells in the peripheral blood correlated closely with anti-tumor responses, leading to complete tumor control. Human CD4^+^ T cell levels revealed similar expansion and persistence to CD8^+^ T cells in TIGIT/28 and TIGIT/28F groups, further contributing to tumor eradication (**Fig. 4D**). Peripheral T cell expansion and persistence was associated with prominent CD4^+^ T cell infiltration in the spleen, observed predominantly in TIGIT/28 and TIGIT/28F treated animals (**Suppl. -Fig. 4B**). Infiltrating T cells remained TIGIT^high^. Human T cell infiltration was not detected in the tumors of non-responding mice, underscoring the critical role of peripheral T cell expansion and persistence in mediating effective anti-tumor responses.

Collectively, these findings demonstrate that TIGIT/28F CSR has a robust anti-tumor efficacy and leads to persistence of TCR T cells, resulting in a significant survival advantage for treated mice.

### TIGIT/28F is associated with enhanced activation and increased mitochondrial respiration upon TCR-specific T cell stimulation

To evaluate the additive stimulatory effect conferred by the TIGIT/28 CSR and test potential compensatory upregulation of other inhibitory receptors upon tumor recognition, we analyzed the expression of several surface molecules associated with T cell activation. TIGIT/28F TCR T cells or TCR T cells without additional CSR were co-cultured with target tumor cells for 48 h. Following co-culture, the expression levels of CD25, CD27, TIM-3, LAG-3, PD-1 and CD226 were analyzed by flow cytometry. The analysis showed upregulation of CD25 in TIGIT/28F and control TCR T cells, yet TIGIT/28F T cells exhibited significantly higher CD25 expression (mean 64.0% vs. 41.7%, **Fig. 5A**). No difference was observed in expression of the co-stimulatory CD27, consistent with similar results for CD28 and CD57 expression. Expression of inhibitory immune checkpoint receptors TIM-3, LAG-3 and PD-1 were also upregulated. Notably, LAG-3 and TIM-3 were higher in TIGIT/28F T cells (mean 52.0% vs. 35.9% and mean 89.4% vs. 82.0%, respectively, **Fig. 5A**). However, PD-1 expression showed no significant difference between the groups. The activating receptor CD226, which competes with TIGIT for binding to CD155 showed high steady-state levels for both groups, yet values were even higher in TIGIT/28F compared to control T cells (**Fig. 5A**).

**Figure 5:**
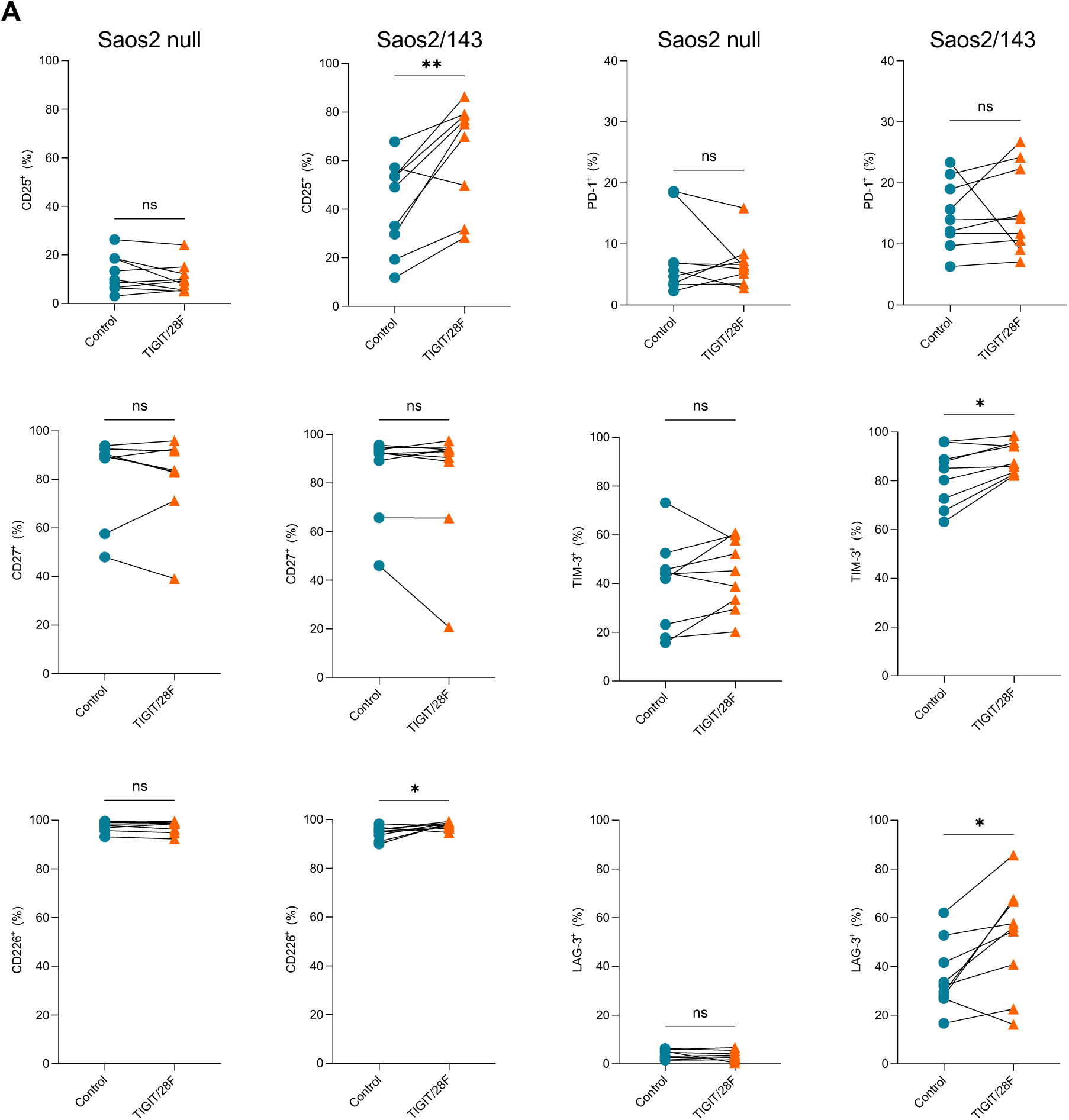
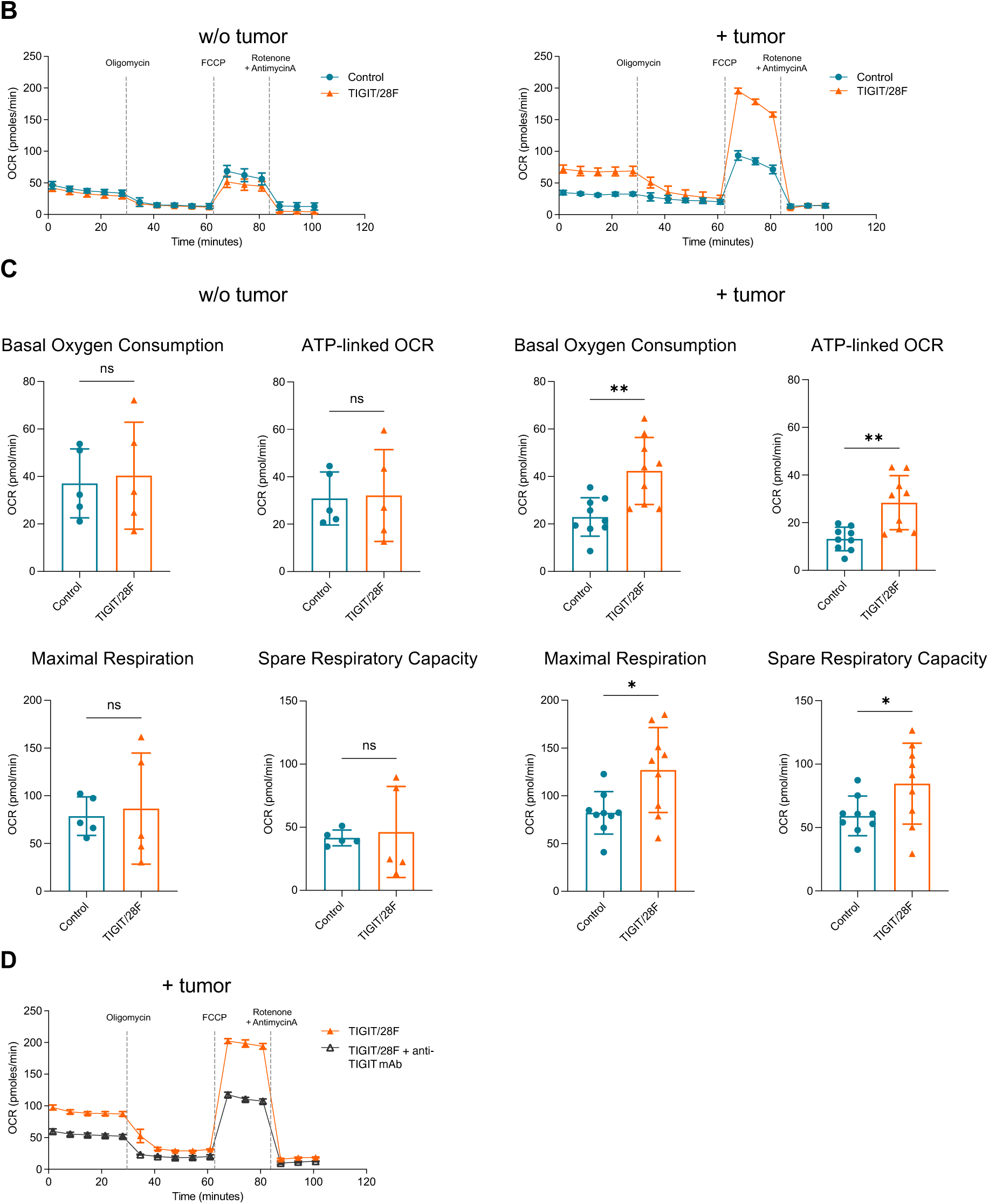
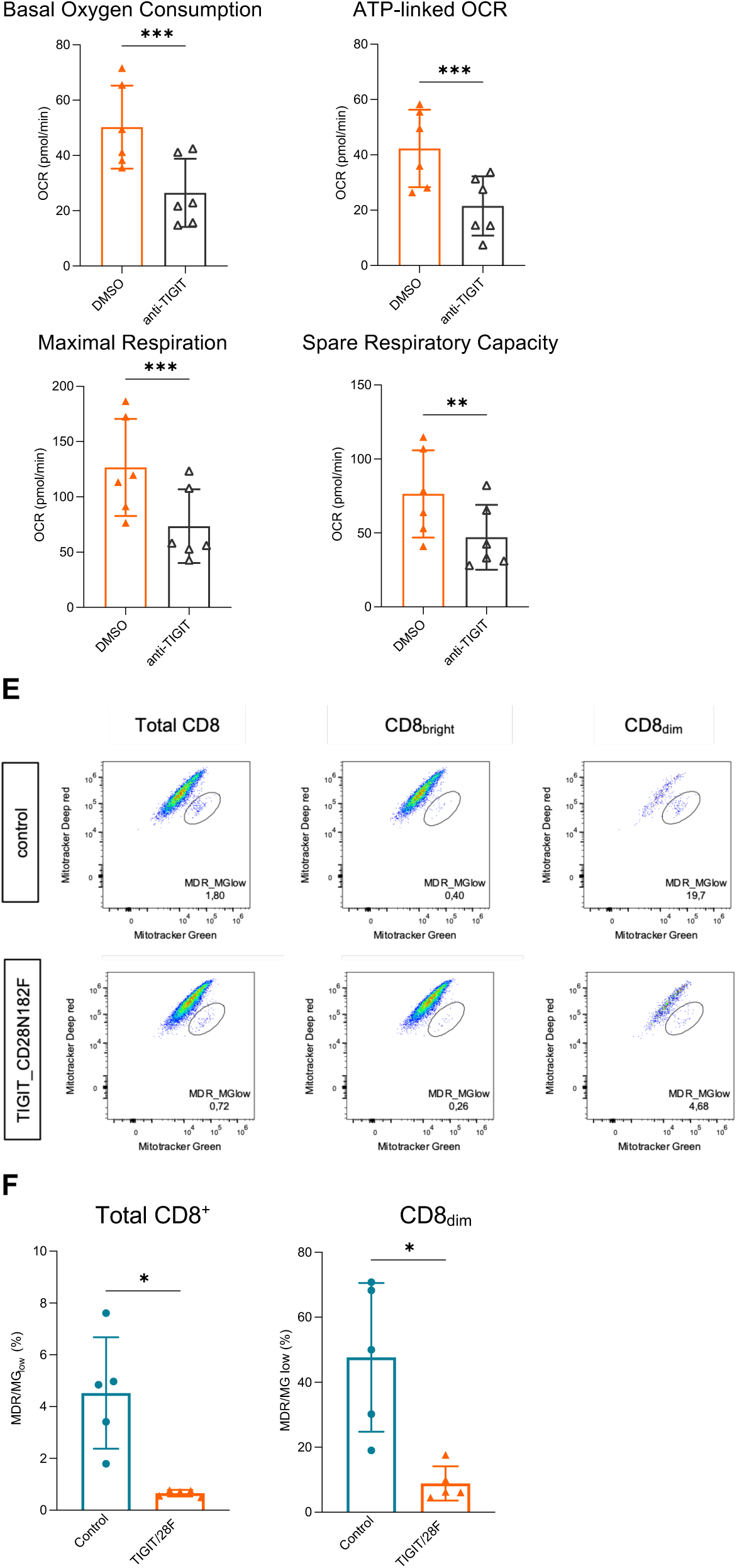
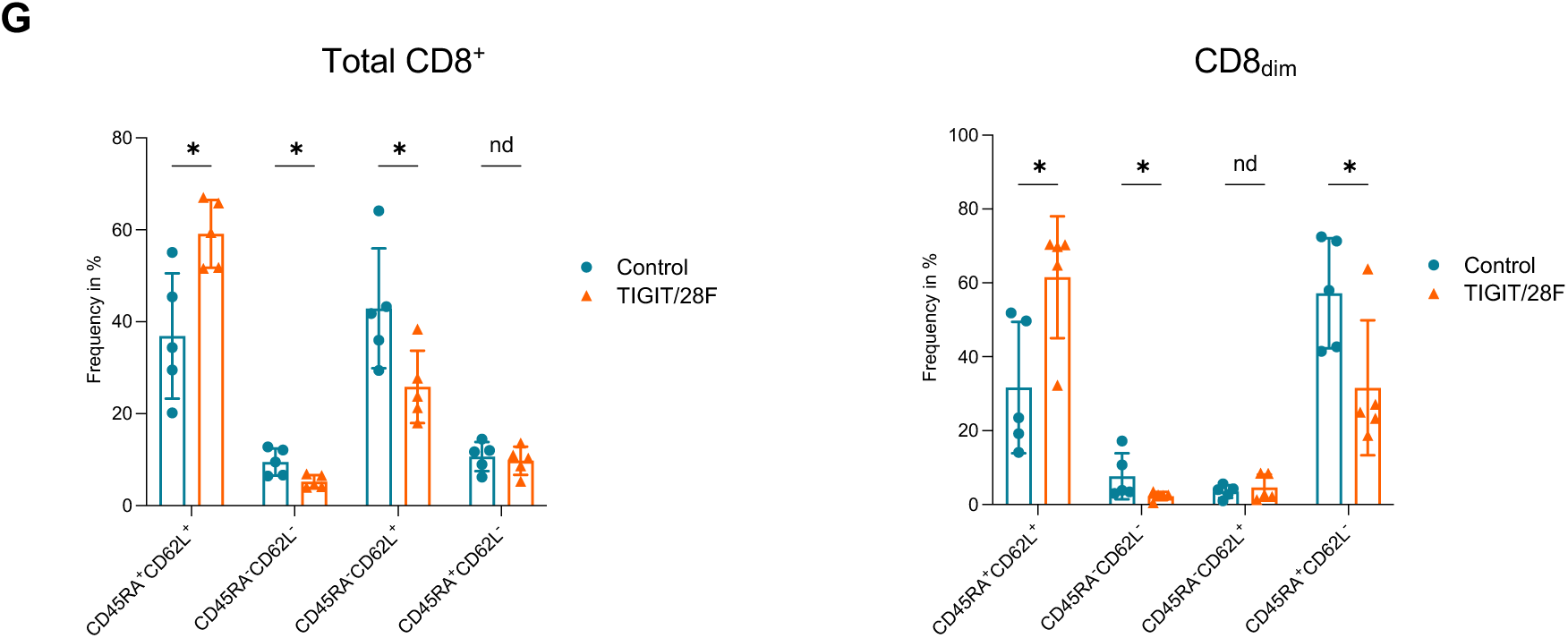
Assessment of the metabolic changes in TIGIT/28F CSR T cells. **(A)** Surface expression of indicated (activation or inhibitory immune checkpoint) receptors analyzed by flow cytometry after 48h co-culture with Saos2/null (antigen negative target) and Saos2/143 tumor cells (CD25: *p*=0.2546 and *p*=0.0029, CD27: *p*=0.6189 and *p*=0.4010, CD226: *p*=0.4545 and *p*=0.0316, PD-1: *p*=0.5754 and *p*=0.7197, TIM-3: *p*=0.2508 and *p*=0.0108, LAG-3: *p*=0.5978 and *p*=0.0178, n=9 biological replicates). **(B)** Example of Seahorse extracellular flux ‘Mito stress test’ of control and TIGIT/28F T cells after stimulation with anti-CD3/CD28 beads and IL-2 (without tumor) and after 48h co-culture with Saos2/143. **(C)** Cumulative data of basal (*p*=0.6940 and *p*=0.0026) and ATP-linked (*p*=0.8593 and *p*=0.0021) oxygen consumption rate (OCR), maximal respiration (*p*=0.7539 and *p*=0.0104) and spare respiratory capacity (*p*=0.7865 and *p*=0.0297) calculated from B (n=5 biological replicates). **(D)** Example of Seahorse extracellular flux ‘Mito stress test’ and cumulative data of basal (*p*=0.0001) and ATP-linked (*p*=0.0001) oxygen consumption rate (OCR), maximal respiration (*p*=0.0008) and spare respiratory capacity (*p*=0.0028) of TIGIT/28F T cells +/- anti-TIGIT blocking mAb (n=6 biological replicates). **(E)** Example of flow cytometric MitoTracker Green and MitoTracker Deep Red measurement in CD8^+^ T cells after co-culture with Saos2/143. **(F)** Frequency of MDR/MG_low_ in total CD8 (*p*=0.0184) and CD8_dim_ T cells (*p*=0.0162, n=5 biological replicates). **(G)** Frequency of naïve (*p*=0.0279), effector (*p*=0.1109), central memory (*p*=0.3869) and EMRA T cells (*p*=0.0281) in control and TIGIT/28F T cells based on the surface expression of CD45RA and CD62L (n=5 biological replicates). Statistical significance in A, C, D and F was determined by two-tailed Student’s *t*-test. * p < 0.05; ** p < 0.01; *** *p* < 0.001; n.s. = not significant. In G multiple paired *t*-test were performed, with the False Discovery Rate approach to correct for multiple comparisons (Q<10%). An asterisk represents a “discovery” (q < Q), while “nd” represents “not a discovery” (q ≥ Q).

Disturbed mitochondrial function after persistent antigen exposure has been linked to terminal exhaustion and impaired self-renewal of T cells ^34–36^. To further elucidate the mechanisms by which the TIGIT/28F CSR enhances anti-tumor responses, we investigated the metabolic activity of the TCR T cells. Similar to the previous experiment, TIGIT/28F TCR T cells or control TCR T cells without additional CSR were co-cultured with target tumor cells for 48 h. Metabolic profiling of T cells was performed using the Seahorse extracellular flux assay. Interestingly, TIGIT/28F TCR T cells demonstrated a marked increase in basal oxygen consumption rate (OCR), a key indicator of mitochondrial respiration, upon TCR-specific stimulation compared to their TIGIT/28F TCR T cells cultured without tumor cells (mean OCR: 23.0 vs. 42.4 pmol/min, **Fig. 5C**). This observation suggests heightened metabolic activation in response to tumor-specific engagement. In contrast, extracellular acidification rate (ECAR), a marker of glycolytic activity, showed no statistically significant difference between the groups, in both steady-state condition and upon antigen-specific stimulation by the co-culture with target tumor cells (**Suppl. -Fig. 5A**).

These results are consistent with the upregulation of CD25 and inhibitory receptors and underscore the role of TIGIT/28F CSR in driving metabolic adaptations that support enhanced anti-tumor activity. Interestingly, the differences observed in mitochondrial parameters between TIGIT/28F and control T cells occurred upon antigen-specific stimulation by the co-culture with target tumor cells, but not in T cells stimulated without tumor cells (**Fig. 5B-C**).

To confirm that the enhanced mitochondrial activity, as indicated by increased oxygen consumption, was mediated by the CSR, Seahorse extracellular flux assays were conducted in the presence of TIGIT blocking antibody. Addition of anti-TIGIT mAb resulted in a significantly reduced basal OCR (26.5 vs. 50.3 pmol/min) and spare respiratory capacity (47.1 vs. 76.4 pmol/min, **Fig. 5D**). No metabolic differences were observed in control T cells treated with the same anti-TIGIT mAb (**Suppl. -Fig. 5B**), indicating that the observed metabolic alterations were primarily driven by additional co-stimulation through the CSR rather than the prevention of inhibitory TIGIT signaling. Consistently, anti-TIGIT blockade reduced the glycolytic activity (as measured by ECAR values) of TIGIT/28F T cells but not control T cells upon antigen-specific stimulation by the co-culture with target tumor cells (**Suppl. - Fig. 5C**).

To further investigate mitochondrial features, mitochondrial mass and membrane potential were assessed by MitoTracker Green (MG) and MitoTracker Deep Red (MDR) co-staining (**Fig. 5E**). TIGIT/28F TCR T cells or control TCR T cells were co-cultured with target tumor cells for two consecutive rounds. Following co-culture, T cells were stained with MDR and MG along with CD8, CD45RA and CD62L surface markers. CD8^+^ TIGIT/28F TCR T cells exhibited no significant differences in mitochondrial membrane potential and showed a moderate increase in mitochondrial mass as compared to control TCR T cells (**Suppl. -Fig. 5D**). Notably, the frequency of T cells with reduced MDR to MG ratio (MDR/MG_low_ population, **Fig. 5E**) was lower in TIGIT/28F TCR T cells compared to control TCR T cells (0.65% vs. 4.52%, **Fig. 5F**), indicating a reduced fraction of T cells with lower mitochondrial activity per mass, and therefore a lower number of cells with metabolic “dysfunction”. Consistently, high frequency of MDR/MG_low_ population is associated with terminal exhaustion in tumor infiltrating lymphocytes ^44^. Interestingly, within the CD8 population, a subgroup of T cells with reduced CD8 expression (CD8_dim_) displayed an abundant MDR/MG_low_ population (**Fig. 5F**). Control CD8_dim_ TCR T cells exhibited a higher frequency of the MDR/MG_low_ population (47.7% vs. 8.9%, **Fig. 5F**). Despite comparable frequencies of CD8_dim_ and CD8_bright_ T cells between TIGIT/28F and control TCR T cells (**Suppl. -Fig. 5E**), their cellular phenotype differed markedly. Both control and TIGIT/28F CD8_dim_ TCR

T cells expressed CD45RA as marker of quiescence. However, control TCR T cells were predominantly composed of terminally differentiated T_EMRA_ (CD45RA^+^/CD62L^-^), while TIGIT/28F TCR T cells were primarily composed of less-differentiated naïve T cells (CD45RA^+^/CD62L^+^, **Fig. 5G**). In the total CD8 population, TIGIT/28F TCR T cells exhibited a more naïve phenotype compared to the central memory-dominated phenotype (T_CM_, CD45RA^-^/CD62L^+^) of control TCR T cells.

Taken together, TIGIT/28F TCR T cells exhibited enhanced mitochondrial activity, a lower frequency of T cells with an exhaustion-associated mitochondrial phenotype and a preserved less-differentiated, quiescent phenotype compared to control TCR T cells when co-cultured with target tumor cells. This cellular and metabolic “fitness” may account for the improved persistence and anti-tumor responses observed in the xenograft tumor model.

### TIGIT/28F CSR improves cytokine response and cytotoxicity of CAR T cells

To extend the applicability of the TIGIT/28F CSR beyond TCR-gene therapy against solid tumor, we assessed its functional potential in the OCI-AML3 acute myeloid leukemia (AML) model. This leukemic cell line expresses the p53TCR-restricted antigen and the CSR ligand CD155, albeit at a lower MFI compared to Saos2/143 (**Fig. 6A**). For the cytotoxicity assay, OCI-AML3 cells expressing firefly luciferase (Fluc) were cultured with p53TCR-T cells co-expressing TIGIT/28F or without CSR (control T cells) for 24 h and specific lysis was determined via luciferase assay. Control T cells showed a mean tumor cell lysis of 64.9%. In contrast, TIGIT/28F TCR T cells (mean 82.3%) demonstrated significantly enhanced tumor cell lysis compared to the control TCR T cells (**Fig. 6B**).

**Figure 6:**
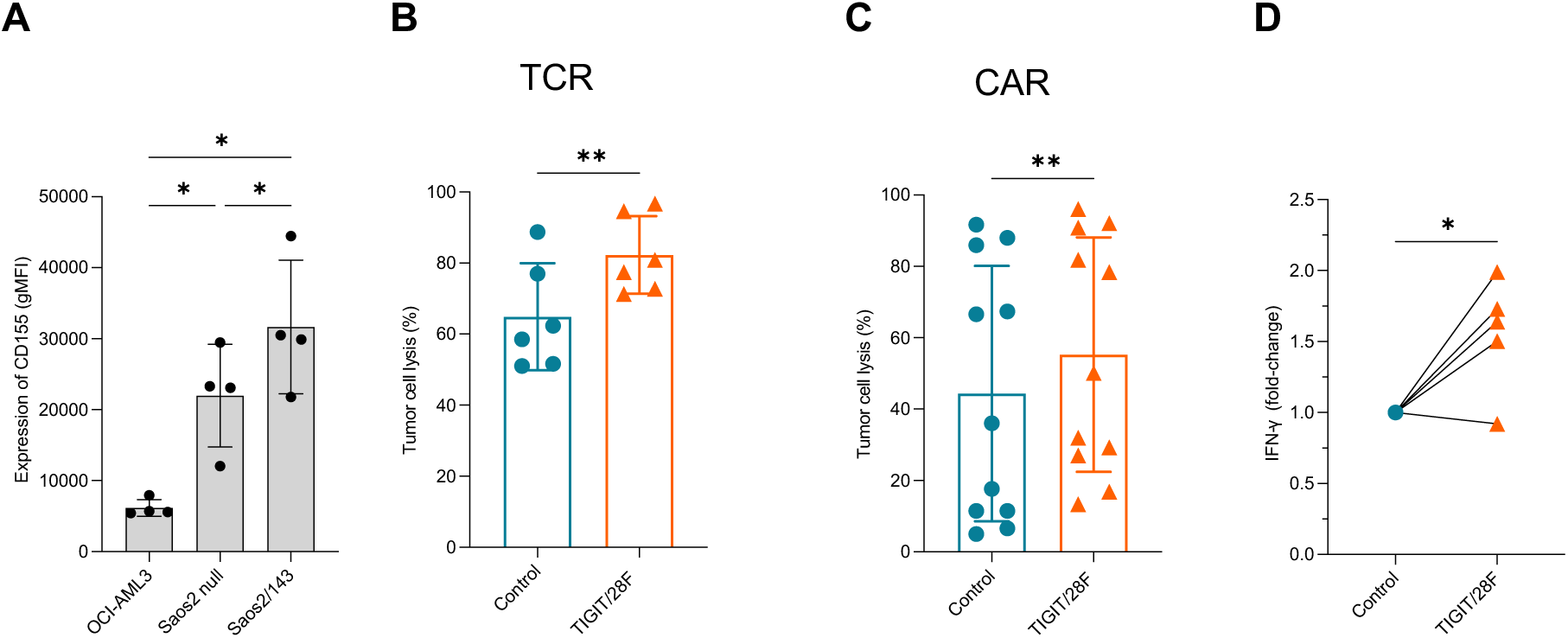
Functional characteristics of the TIGIT/28F CSR in CAR-T cells. **(A)** Expression of CD155 of different target tumor cell lines indicated by gMFI of flow cytometric measurement (OCI vs. Saos2 null *p*=0.0345, OCI vs. Saos2/143 *p*=0.0183 and Saos2 null vs. Saos2/143 *p*=0.0294 n=4). **(B)** and **(C)** Lysis of OCI-AML3 tumor cells after 24 h co-culture with T cells transduced with tumor antigen-specific TCR **(B)** or CAR **(C)** in combination with TIGIT/28F CSR or control (*p*=0.0049 and n=6 in (B) and *p*=0.0059 and n=11 biological replicates in (C)). **(D)** Release of IFN-γ by NKp30 CAR-T cells as fold-change after co-culture with OCI-AML3, measured by ELISA (*p*=0.0352, n=5 biological replicates). Statistical significance in A was determined by one-way ANOVA and Tukey’s multiple comparisons test and in B-D by by two-tailed Student’s *t*-test. * *p* < 0.05; ** *p* < 0.01; *** *p* < 0.001; n.s. = not significant.

We also assessed the functional potential of the optimized CSR in CAR-engineered T cells using the same OCI-AML3 target cell line. T cells were equipped with a 4-1BB-based second-generation NKp30-CAR ^37^ targeting B7-H6 (natural killer cell cytotoxicity receptor 3 ligand 1, NCR3LG1). TIGIT/28F or control (without CSR) CAR T cells were cultured with B7-H6 expressing OCI-AML3 cells (**Suppl. - Fig. 6**). TIGIT/28F CAR T cells exhibited a stronger cytolytic response compared to control CAR T cells (**Fig. 6C**). The levels of tumor-induced secretion of IFN-γ by CSR-expressing CAR T cells were increased compared to control CAR T cells (1.6 mean fold-change, **Fig. 6D**).

## Discussion

Despite major advances in T cell-based immunotherapies, efficacy in solid cancer remains limited by the expression of suppressive immune checkpoints within the tumor microenvironment, which drive T cell exhaustion and tumor evasion. Immune checkpoint inhibitors have been developed and widely used in cancer immunotherapy, with anti-CTLA-4 and anti-PD1 being now part of standard therapy for several advanced cancers. Despite high numbers of successful immunotherapy trials with ICI, many patients do not respond or show a short-lived response. Key challenges remain the resistance and immune-related adverse events to ICI and ultimately cancer relapse ^38^.

Engineering anti-tumor T cells with immune checkpoint-based CSRs could circumvent these limitations and offer additional therapeutic tools to improve T cell-based cancer immunotherapy. Over the last few years, several CSRs targeting different inhibitory immune checkpoints have been generated and pre-clinically tested ^39^. One of the earliest CSR constructs fused the extracellular domain of PD-1 with the intracellular signalling domain of CD28, thereby demonstrating the feasibility of using CSRs to convert inhibitory signals into co-stimulatory cues and enhance T cell activation ^40^. Other studies showed that replacing the CD28 transmembrane (TM) domain with the corresponding domain of PD-1 further improved functionality of murine TCR-T cells ^41^, or exchanging CD28 with other co-stimulatory domains, such as 4-1BB, enhanced efficacy of CAR-T cells in xenograft metastatic tumor models ^42^. In addition to PD-1-based constructs, CSRs have been designed using extracellular and TM domains derived from other inhibitory receptors, including CTLA-4 ^43,44^ or TIM-3 ^16,45^, highlighting the importance of receptor architecture for optimal signalling and function of antigen-receptor modified T cells.

In this study, we designed a CSR targeting the emerging checkpoint TIGIT ^46^. To date, only two TIGIT CSRs have been described: TIGIT-CD28 ^17^ and PD-1/TIGIT-CD28 ^47^. Here, we designed two different TIGIT/28-based CSRs and evaluated their functional properties in TCR- but also CAR-engineered human T cells.

While our first (original) TIGIT/28 CSR design improves the cellular fitness of TCR-modified human T cells *in vitro*, the therapeutic benefit in xenograft tumor model was rather modest, mainly due to premature T cell differentiation induced by the CSR and therefore a short persistence of anti-tumor T cells *in vivo*. Premature exhaustion of TCR/CAR gene-modified T cells is a hallmark of tumor evasion and is often associated with poor therapeutic outcomes in cancer patients. T cell persistence was shown to be critical for long-term tumor control in tumor-infiltrating lymphocyte (TIL) and CAR T cell therapy studies ^48,49^. In patients treated with TCR T cells, T cell persistence was reported to be associated with a less-differentiated T cell phenotype, including preserved expression of CD28 and more effective anti-tumor responses ^50,51^. Studies have clearly demonstrated that the type of costimulatory domain used in CAR constructs influences T cell persistence ^52,53^, showing that CAR T cells with CD28-based co-stimulation exhibit rapid expansion but shorter persistence compared to 4-1BB stimulated CAR T cells ^54,55^. This poorer persistence and premature exhaustion of T cells expressing a CD28-based second-generation CAR were linked to excessive NFAT production by the YMNM motif in the CD28 signaling domain ^33,56^. This signaling motif is shared by ICOS, another co-stimulatory receptor of the B7-CD28 family. These motifs differ in only one amino acid, namely phenylalanine in ICOS (YMFM) vs. asparagine in CD28 (YMNM). Substituting the asparagine (N) in CD28 by phenylalanine (F) in a second-generation CAR resulted in improved anti-tumor responses, less T cell exhaustion and differentiation^33^. Using site-directed mutagenesis, we introduced this amino acid change (N to F) in the TIGIT/28 CSR to modulate the activation signal of the CSR and prevent early differentiation.

Expression of the modified TIGIT/28F (optimized) CSR in TCR-transduced T cells demonstrated improved cytotoxicity against tumor cells *in vitro*, comparable to the original CSR, while maintaining proliferation and a less-differentiated phenotype. In sharp contrast to the original CSR, expression of the optimized CSR resulted in long-term persistence of infused TCR T cells and complete tumor eradication in xenograft tumor model. To date, many FDA approved CARs and those tested in clinical trials are CD28-based constructs, raising the question of excessive CD28-signaling mediated short-term persistence and thus the risk of patient relapse. Notably, CAR T cells utilizing the full transmembrane domain of ICOS appeared less cytotoxic ^33^. A simple single amino acid change in the CD28 signaling domain therefore helped to substantially improve the *in vivo* persistence, while maintaining the potent cytolytic response of CD28-costimulated T cells. Furthermore, differences in co-stimulatory signaling were reported to drive T cell metabolism from glycolysis towards mitochondrial respiration, accompanied by central memory formation and enhanced *in vivo* persistence ^52^. Modified co-stimulation provided by the optimized CSR may have influenced the metabolic activity of TCR T cells in the context of repetitive tumor challenges, resulting in a less-differentiated phenotype and better persistence.

Combining CSR with T cell-based therapies bears great potential to enhance their efficacy. However, CSR can also lead to potential off-target toxicities against non-malignant cells ^57^, raising a safety concern ^39^ in clinical translation. In our study, the optimized CSR showed no reactivity against target cells in the absence of an antigen-specific TCR. Therefore, the careful design of CSR co-stimulation offers the possibility to tune the safety profile of T cells, avoid tumor-induced premature differentiation and further improve anti-tumor responses. Although no CSR has yet received regulatory approval for clinical use, several CSRs are currently being evaluated in TCR/CAR-T cells in early-phase clinical trials in patients with recurrent glioblastoma ^58^, metastatic malignancies ^59^ or in patients with relapsed/refractory B cell lymphomas, with one study reporting a complete remission rate of 41.2% among 17 treated patients, without the occurrence of severe cytokine release syndrome or neurologic toxicity ^60^.

Collectively, these clinical studies provide initial evidence supporting the feasibility, safety, and potential therapeutic activity of CSR-engineered T cells, although validation in larger patient cohorts is still needed. In this context, our novel optimized TIGIT CSR offers promising therapeutic perspectives to further improve immune T cell fitness for effective immunotherapy, in particular for solid, hard-to-treat cancer with hostile TME, where the transfer of modified T cells has so far only achieved limited success.

## Materials and Methods

### Cloning of TIGIT CSR constructs

The construct for TIGIT/28 was designed via SnapGene (Dotmatics) and is composed of the signal peptide (amino acid 1-21) and extracellular domain (amino acid 22-141) of TIGIT (NCBI ref. NM_173799.3) and the transmembrane (amino acid 153-179) and cytoplasmic domain (amino acid 180-220) of CD28 (NCBI ref. NM_006139.4). DNA synthesis was performed by Invitrogen GeneArt (ThermoFischer Scientific, Germany). The construct was cloned into a pMX-IRES-Puro (RTV014) retroviral vector backbone using BamHI/NotI restriction enzyme sites.

TIGIT/28F was generated from TIGIT/28 by introducing an amino acid change from asparagine to phenylalanine at position 182. Site directed mutagenesis was performed via PCR using Q5® High-Fidelity DNA Polymerase (NEB, Ipswich, USA) with primers displayed in **Suppl.-Table 1**. DpNI was added to the PCR products to degrade methylated template DNA. PCR products were transformed into competent bacterial strain JM109 via heat-shock and DNA was purified EndoFree Plasmid Maxi Kit (QIAGEN, Hilden, Germany). Plasmid sequencing was performed by Eurofins MWG Operon (Luxembourg) and results were aligned using SnapGene (Dotmatics).

### Human T cell culture and tumor cell lines

Human peripheral blood mononuclear cells (PBMCs) were isolated from buffy coats via density gradient centrifugation with Histopaque-1077 (Sigma-Aldrich, St. Louis, USA). Buffy coats were obtained from HLA_A2^-^ healthy donors and were provided by the Transfusion Center of the University Medical Center Mainz. *In vitro*, PBMCs/T cells were cultured in RPMI-1640 supplemented with 10% heat-inactivated human AB-Serum, 2 mM L-Glutamine, 25 mM 2-(4-(2-Hydroxyethyl)-1-piperazinyl)-ethansulfonacid (HEPES), 100 U/ml penicillin and 0.1 mg/ml streptomycin (huRPMI). As described before ^29,30^, T cells were activated once a week by the addition of 5 µl Dynabeads Human T Activator CD3/CD28 (Gibco, Carlsbad, USA) per 1 × 10^6^ cells and 600 U/ml human IL-2 (Proleukin S, Novartis, Basel, Switzerland). Proliferation *in vitro* was assessed by calculating the population doubling levels (PDL) with the following formula:

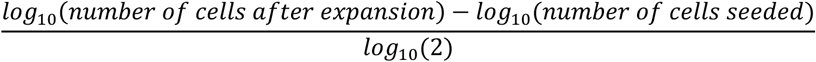

The HLA_A2.1^+^ p53 null osteosarcoma cell line Saos2 and its p53-transfectant Saos2/143 were cultured in RPMI-1640 supplemented with 10% heat-inactivated FCS. The acute myeloid leukemia cell lines OCI-AML3 and OCI-AML3 stably expressing firefly luciferase and EGFP were cultured in RPMI-1640 supplemented with 10% fetal calf serum (FCS, Sigma Aldrich), 100 U/ml penicillin and 0.1 mg/ml streptomycin (P/S, Sigma), and 1% L-Glutamine (L-Glu, Sigma).

### Retroviral Transduction of human PBMCs/T cells

Tumor-antigen specific TCR (scp53TCR) or CAR (NKp30 CAR) and the chimeric switch receptors (TIGIT/28 and TIGIT/28F) were stably integrated into human T cells via retroviral transduction as described previously ^30,61^. Briefly, the packaging cell line Phoenix-AMPHO was transfected with 10 µg transgene plasmid DNA and 5 µg helper plasmids (pHit60 and pColt-Galv) using transfectant reagent Fugene 6 (Promega, Madison, USA) to produce virus particles. Human PBMCs stimulated with 30 ng/ml soluble anti-CD3 antibody (OKT-3) and 600 U/ml IL-2 for 48 h were resuspended in retroviral supernatant in 24 well/plate. To increase transduction efficiency, 5 µg/ml hexadimethrine bromide (Polybrene, Sigma-Aldrich, St. Louis, USA) was added. Cells were centrifuged at 600xg, 32°C for 90 min and subsequently incubated at 37°C overnight. The viral supernatant was removed, and T cells were re-stimulated as described above and selected for gene expression by addition of puromycin (5 µg/ml, overnight on day 6 post transduction) and/or neomycin (800 µg/ml for 7 days).

### Flow Cytometry

For cell surface staining, T cells were stained using indicated antibodies for 15 min at room temperature. A list of antibodies used for flow cytometry is provided in **Suppl.-Table 2**.

Secretion of lytic granules (degranulation assay) upon co-culture with target tumor cells was determined by measuring the expression of LAMP1 (CD107a) on the cell surface. T cells were activated and incubated for 24 h in the presence of monensin (Monensin Solution 1000X, eBioscience, San Diego, USA) to block Golgi transport and the CD107a antibody. After incubation, T cells were washed with PBS and stained for extracellular antigens as described above.

Fluorescent MitoTracker Deep Red and Green (Cell Signaling Technology, Cambridge, United Kingdom) were used for analysis of mitochondrial mass and membrane potential. After washing with PBS, T cells were incubated in RPMI 1640 containing 10 nM Mitotracker DeepRed and 50 nM MitoTracker Green for 30 min at 37°C, 5% CO_2_. Reaction was stopped by washing with cold (4°C) PBS and the cells were stained for surface markers as described above.

Flow cytometric measurement was performed on a FACS Canto II (BD, Franklin Lakes, USA) and MitoTracker measurements on an Aurora spectral flow cytometer (Cytek Biosciences, Fremont, USA). FlowJo V10.8 software (BD, Franklin Lakes, USA) was used for data analysis.

### *In vitro* Cytotoxicity Assays

*<u>Short-term T cell cytotoxicity</u>* was evaluated by performing standard ^51^Cr-release assays ^62^. In brief, target tumor cells were labelled with 100 µCi radioactive sodium ^51^chromate (Na_251_CrO_4_) for 90 min at 37 °C. Then, target cells were washed with huRPMI and co-cultured with T cells at different E:T ratios for 5 h in 200 µl huRPMI. Chromium release was measured with a gamma counter (Canberra Packard Cobra II, Perkin Elmer) in 100 µl cell-free supernatant of the co-culture. Spontaneous lysis and maximum release were determined in cell-free supernatant from tumor cells without T cell co-culture and medium containing labelled tumor cells, respectively. Specific lysis was finally calculated using the following formula:

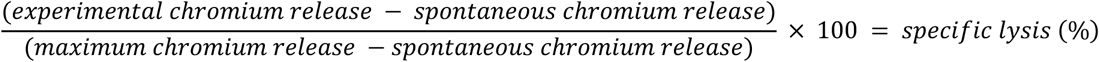

*<u>Long-term T cell cytotoxicity</u>* was assessed in tumor colony-forming assays. T cells were co-cultured with the target antigen expressing tumor cell line Saos2/143 or non-target Saos2 cell line. If not otherwise specified, tumor cells were seeded in a 6-well plate and the corresponding number of T cells (indicated by the E:T ratio) was added. T cells and tumor cells were co-cultured for 24 h at 37°C and 5% CO_2_. Then, the supernatant containing T cells was transferred to a new well with freshly seeded tumor cells for another round of co-culture. If not otherwise indicated, three rounds of co-culture (24 h each) were performed. Remaining viable tumor cells were washed with PBS and fixed with 4% PFA for 10 min and stained by adding 400 µl of 1% crystal violet dye (Sigma-Aldrich, St. Louis, USA) for 15 min at room temperature. The dye was removed and plates washed with PBS. Dye absorption was analysed by adding 500 µl of 5% SDS in PBS (to dissolve the dye) and quantified by measuring the OD_570_ in a multimode microplate reader (Varioskan LUX, ThermoFischer Scientific, Waltham, USA). For blockade of TIGIT signaling, a monoclonal anti-TIGIT blocking antibody was used at a concentration of 4 µg/ml (MBSA43, eBioscience, Invitrogen, Waltham, USA).

### Luciferase-based Cytotoxicity Assay

The amount of living target cells (OCI-AML3) after co-culture with effector cells was determined based on the light emission of the intracellular luciferase activity. For the co-culture, target cells were seeded in 100 µl culture medium per well in a white 96-well plate with transparent bottom. Effector cells were added in 100 µl culture medium according to the indicated E:T ratio, using triplicates for each experimental condition. Wells without effector cells were filled up with 100 µl culture medium to assess spontaneous lysis or with 100 µl culture medium supplemented with 0.15 mg/ml Digitonin to assess maximal lysis. D-luciferin (Perkin Elmer, Waltham, USA) was added to each well at a concentration of 0.15 mg/ml. Empty wells were filled with culture medium (blank wells). The co-culture was incubated at 37°C, 5% CO_2_ for 24 h. Light emission from the enzymatic oxidation of D-luciferin in viable target cells was quantified with a multimode microplate reader (Varioskan LUX, Thermo Fischer Scientific, Waltham, USA) after 16 h, 20 h, and 24 h. Specific lysis of target cells was finally calculated with the following formula:

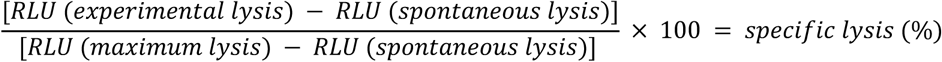

### Enzyme-linked Immunosorbent Assay (ELISA)

Secretion of IFN-γ *in vitro* or *in vivo* was quantified by ELISA (Invitrogen, Carlsbad, USA). The assay was performed according to the manufacturer’s protocol. The OD_450_ was determined with a multimode microplate reader (Varioskan LUX, Thermo Fischer Scientific, Waltham, USA).

### Extracellular flux assays

The Seahorse XFp Analyzer (Agilent) was used for the quantification of the oxygen consumption rate (OCR) and the extracellular acidification rate (ECAR) of TCR-/CSR-transduced T cells and control cells. Before the measurement, T cells were stimulated with anti-CD3/anti-CD28 beads and IL-2 for 3-4 days as described before or by two rounds of 24 h co-culture with target tumor cells at an E:T of 1:1. Blockade of TIGIT signaling was performed by addition of an anti-TIGIT blocking mAb (MBSA43, eBioscience, Invitrogen, Waltham, USA) at a final concentration of 4 µg/ml. To adhere T cells to Culture Miniplates, each well was coated with 50 µl of Poly-D-Lysine (Sigma P6407) for 1 h at room temperature. XFp Sensor Cartridges were hydrated by the addition of 200 µl XF calibrant to each well and 400 µL to each outside well and stored at 37°C in a non-CO_2_ incubator overnight. The Seahorse XFp Cell Mito Stress Test (Agilent) was performed according to the manufacturer’s protocol. T cells were seeded in XFp RPMI Medium (supplemented with 10 mM glucose, 1 mM sodium pyruvate and 2 mM L-Glutamine). Two measurement cycles were added during the baseline phase to achieve stable baseline conditions. Oligomycin, fluoro-carbonyl cyanide phenylhydrazone (FCCP), Rotenone and Antimycin A (Sigma-Aldrich, Burlington, USA) were injected with a final concentration of 1.5 µM, 2 µM and 0.5 µM, respectively. Seahorse Wave Desktop Software was used for further analysis and quality control.

### Animal models

Mice were obtained from the central animal facility of the Johannes Gutenberg University Mainz, Germany. Mice were kept according to the guidelines for animal care of the Johannes Gutenberg University Mainz and all procedures were performed according to the German federal and state regulations and approved by the responsible national authority (National Investigation Office Rhineland-Palatinate).

To study T cell function and anti-tumor cytotoxicity *in vivo*, a xenograft tumor model was employed as described previously ^29,30^. Immunocompromised NOD.Cg-Prkdc^scid^ Il2rg^tm1Wjl^/SzJ (NSG) mice were subcutaneously injected with 3×10^6^ Saos2/143 tumor cells in the left flank (at day 0). After tumor engraftment, 5×10^6^ T cells were injected intravenously in combination with IL-2 (7.2×10^5^ IU) injection (intraperitoneal) at day 5 or 7. Tumor growth was monitored by calculating tumor volume with the following formula: length × width^2^. Mice harboring a tumor ≥ 1cm^3^ were sacrificed. Spleen and tumor of sacrificed animals were used to isolate infiltrating T cells and subsequent flow cytometric analysis as described before ^30^. Blood was drawn at the indicated time points to measure T cell counts. Red blood cells were lysed with 2 ml lysis buffer (NH_4_CL 174 mM, KHCO_3_ 10 mM, Na_2_EDTA 0.1 mM in 500 ml distilled water) and remaining PBMCs were stained for surface markers as described above. Stained PBMCs were washed with PBS, resuspended in 200 µl of 1% PFA solution and 10 µl of Sulfate Latex Beads (4% w/v 5 µm, Invitrogen, Carlsbad, USA) were added. Absolute T cell counts were calculated with the following formula:

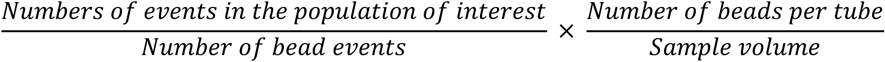

### Statistical analysis and software

Statistical evaluation and graphical reports were established with Graphpad Prism version 10.0.0 (GraphPad Software, San Diego, California, USA). To test for significant differences between samples, repeated measures one-way analysis of variance (one-way ANOVA) or paired *t*-test was performed, followed by Tukey’s multiple comparisons test, as indicated. If not otherwise indicated, results were reported as adjusted p values. Samples of each donor were matched, and Gaussian distribution was tested with the Kolmogorov-Smirnov and Shapiro-Wilk test as well as graphically using the respective Normal Quantile-Quantile (Q-Q) Plot. All survival curves from one experiment were first compared with Log-rank (Mantel-Cox) test. Then, pairs of groups were compared again with the Log-rank (Mantel-Cox) test and Bonferroni-corrected for multiple comparisons.

## Data availability

The data generated in this study are available upon request from the corresponding author.

## Acknowledgments

The authors thank Dr. Catherine Wölfel and Dr. Udo Hartwig (UMC Mainz) for providing the luciferase/GFP-transduced OCI-AML3 cell line and the NKp30 CAR construct; healthy volunteers who donating blood for T cell transduction, the Translational Animal Research Center (TARC) of the UMC Mainz, for breeding and providing the animals.

## Author contributions

H.E. conceived the project, designed and supervised the study. K.J.L. and E.A.F. cloned the chimeric switch receptors (CSRs), performed functional testing of the CSRs in TCR-T cells *in vitro* and *in vivo* xenograft model. U.R.S. performed functional testing of the CSRs in CAR-T cells *in vitro*. E.A.F. and U.R.S. performed metabolic flux assays and A.R. performed mitochondrial tracker flow analysis. U.R.S. and A.R. supported flow cytometry analysis. H.E., K.J.L. and E.A.F. analyzed and interpreted the data, wrote and revised the manuscript. U.R.S. and A.R. edited the manuscript. M.T. analyzed and interpreted the data and revised the manuscript.

## References

1. Baulu, E., Gardet, C., Chuvin, N. & Depil, S. TCR-engineered T cell therapy in solid tumors: State of the art and perspectives. Sci. Adv. 9, eadf3700 (2023).

2. Zugasti, I. et al. CAR-T cell therapy for cancer: current challenges and future directions. Sig Transduct Target Ther 10, 210 (2025).

3. Uslu, U. & June, C. H. Beyond the blood: expanding CAR T cell therapy to solid tumors. Nat Biotechnol 43, 506–515 (2025).

4. Crespo, J., Sun, H., Welling, T. H., Tian, Z. & Zou, W. T cell anergy, exhaustion, senescence, and stemness in the tumor microenvironment. Current Opinion in Immunology 25, 214–221 (2013).

5. Engelhardt, J. J., Sullivan, T. J. & Allison, J. P. CTLA-4 Overexpression Inhibits T Cell Responses through a CD28-B7-Dependent Mechanism. The Journal of Immunology 177, 1052–1061 (2006).

6. Hui, E. et al. T cell costimulatory receptor CD28 is a primary target for PD-1–mediated inhibition. Science 355, 1428–1433 (2017).

7. Chauvin, J.-M. et al. TIGIT and PD-1 impair tumor antigen–specific CD8+ T cells in melanoma patients. J. Clin. Invest. 125, 2046–2058 (2015).

8. Li, X., Shao, C., Shi, Y. & Han, W. Lessons learned from the blockade of immune checkpoints in cancer immunotherapy. J Hematol Oncol 11, 31 (2018).

9. Tay, J. C., Zha, S. & Wang, S. Chimeric Switch Receptor: Switching for Improved Adoptive T-Cell Therapy Against Cancers. Immunotherapy 9, 1339–1349 (2017).

10. Irvine, D. J., Maus, M. V., Mooney, D. J. & Wong, W. W. The future of engineered immune cell therapies. Science 378, 853–858 (2022).

11. June, C. H., O’Connor, R. S., Kawalekar, O. U., Ghassemi, S. & Milone, M. C. CAR T cell immunotherapy for human cancer. Science 359, 1361–1365 (2018).

12. Neelapu, S. S. et al. Axicabtagene Ciloleucel CAR T-Cell Therapy in Refractory Large B-Cell Lymphoma. N Engl J Med 377, 2531–2544 (2017).

13. Schlenker, R. et al. Chimeric PD-1:28 Receptor Upgrades Low-Avidity T cells and Restores Effector Function of Tumor-Infiltrating Lymphocytes for Adoptive Cell Therapy. Cancer Research 77, 3577–3590 (2017).

14. Liu, X. et al. A Chimeric Switch-Receptor Targeting PD1 Augments the Efficacy of Second-Generation CAR T Cells in Advanced Solid Tumors. Cancer Research 76, 1578–1590 (2016).

15. Liang, Y. et al. CD19 CAR-T expressing PD-1/CD28 chimeric switch receptor as a salvage therapy for DLBCL patients treated with different CD19-directed CAR T-cell therapies. J Hematol Oncol 14, 26 (2021).

16. Zhao, S. et al. Switch receptor T3/28 improves long-term persistence and antitumor efficacy of CAR-T cells. J Immunother Cancer 9, (2021).

17. Hoogi, S. et al. A TIGIT-based chimeric co-stimulatory switch receptor improves T-cell anti-tumor function. j. immunotherapy cancer 7, 243 (2019).

18. Worboys, J. D. et al. TIGIT can inhibit T cell activation via ligation-induced nanoclusters, independent of CD226 co-stimulation. Nat Commun 14, 5016 (2023).

19. Hasan, M. F. et al. Knockout of the inhibitory receptor TIGIT enhances the antitumor response of ex vivo expanded NK cells and prevents fratricide with therapeutic Fc-active TIGIT antibodies. J Immunother Cancer 11, e007502 (2023).

20. Xiao, S. et al. Checkpoint Receptor TIGIT Expressed on Tim-1+ B Cells Regulates Tissue Inflammation. Cell Reports 32, 107892 (2020).

21. Hasan, M. M. et al. Implication of TIGIT+ human memory B cells in immune regulation. Nat Commun 12, 1534 (2021).

22. Johnston, R. J. et al. The Immunoreceptor TIGIT Regulates Antitumor and Antiviral CD8 + T Cell Effector Function. Cancer Cell 26, 923–937 (2014).

23. Yu, X. et al. The surface protein TIGIT suppresses T cell activation by promoting the generation of mature immunoregulatory dendritic cells. Nat Immunol 10, 48–57 (2009).

24. Gao, J., Zheng, Q., Xin, N., Wang, W. & Zhao, C. CD155, an onco-immunologic molecule in human tumors. Cancer Science 108, 1934–1938 (2017).

25. Zeng, T. et al. The CD112R/CD112 axis: a breakthrough in cancer immunotherapy. J Exp Clin Cancer Res 40, 285 (2021).

26. He, W. et al. CD155T/TIGIT Signaling Regulates CD8+ T-cell Metabolism and Promotes Tumor Progression in Human Gastric Cancer. Cancer Research 77, 6375–6388 (2017).

27. Guillerey, C. et al. TIGIT immune checkpoint blockade restores CD8+ T-cell immunity against multiple myeloma. Blood 132, 1689–1694 (2018).

28. Harjunpää, H. & Guillerey, C. TIGIT as an emerging immune checkpoint. Clinical and Experimental Immunology 200, 108–119 (2020).

29. Echchannaoui, H. et al. A Potent Tumor-Reactive p53-Specific Single-Chain TCR without On- or Off-Target Autoimmunity In Vivo. Molecular Therapy 27, 261–271 (2019).

30. Legscha, K. J. et al. Δ133p53α enhances metabolic and cellular fitness of TCR-engineered T cells and promotes superior antitumor immunity. J Immunother Cancer 9, e001846 (2021).

31. Lorenzo-Herrero, S., Sordo-Bahamonde, C., Gonzalez, S. & López-Soto, A. CD107a Degranulation Assay to Evaluate Immune Cell Antitumor Activity. in Cancer Immunosurveillance (eds López-Soto, A. & Folgueras, A. R.) vol. 1884 119–130 (Springer New York, New York, NY, 2019).

32. Long, A. H. et al. 4-1BB costimulation ameliorates T cell exhaustion induced by tonic signaling of chimeric antigen receptors. Nat Med 21, 581–590 (2015).

33. Guedan, S. et al. Single residue in CD28-costimulated CAR-T cells limits long-term persistence and antitumor durability. Journal of Clinical Investigation 130, 3087–3097 (2020).

34. Yu, Y.-R. et al. Disturbed mitochondrial dynamics in CD8+ TILs reinforce T cell exhaustion. Nat Immunol 21, 1540–1551 (2020).

35. Vardhana, S. A. et al. Impaired mitochondrial oxidative phosphorylation limits the self-renewal of T cells exposed to persistent antigen. Nat Immunol 21, 1022–1033 (2020).

36. Wu, H. et al. Mitochondrial dysfunction promotes the transition of precursor to terminally exhausted T cells through HIF-1α-mediated glycolytic reprogramming. Nat Commun 14, 6858 (2023).

37. Givi, S. et al. CRISPR/Cas9 TCR-Edited NKp30 CAR T Cells Exhibit Superior Anti-Tumor Immunity to B7H6-Expressing Leukemia and Melanoma. IJMS 26, 8235 (2025).

38. Bagchi, S., Yuan, R. & Engleman, E. G. Immune Checkpoint Inhibitors for the Treatment of Cancer: Clinical Impact and Mechanisms of Response and Resistance. Annu. Rev. Pathol. Mech. Dis. 16, 223–249 (2021).

39. Rane, R. et al. Chimeric switch and inverted cytokine receptors in T cell therapy: reprogramming T cells to overcome immune suppression in the solid tumor microenvironment. Front. Immunol. 16, 1662238 (2025).

40. Prosser, M. E., Brown, C. E., Shami, A. F., Forman, S. J. & Jensen, M. C. Tumor PD-L1 co-stimulates primary human CD8+ cytotoxic T cells modified to express a PD1:CD28 chimeric receptor. Molecular Immunology 51, 263–272 (2012).

41. Kobold, S. et al. Impact of a New Fusion Receptor on PD-1–Mediated Immunosuppression in Adoptive T Cell Therapy. JNCI: Journal of the National Cancer Institute 107, (2015).

42. Ma, Q. et al. A PD-L1-targeting chimeric switch receptor enhances efficacy of CAR-T cell for pleural and peritoneal metastasis. Sig Transduct Target Ther 7, 380 (2022).

43. Shin, J. H. et al. Positive conversion of negative signaling of CTLA4 potentiates antitumor efficacy of adoptive T-cell therapy in murine tumor models. Blood 119, 5678–5687 (2012).

44. Prinz, L. F. et al. An anti-CD19/CTLA-4 switch improves efficacy and selectivity of CAR T cells targeting CD80/86-upregulated DLBCL. Cell Reports Medicine 5, 101421 (2024).

45. Blaeschke, F. et al. Design and Evaluation of TIM-3-CD28 Checkpoint Fusion Proteins to Improve Anti-CD19 CAR T-Cell Function. Front. Immunol. 13, 845499 (2022).

46. Kraehenbuehl, L., Weng, C.-H., Eghbali, S., Wolchok, J. D. & Merghoub, T. Enhancing immunotherapy in cancer by targeting emerging immunomodulatory pathways. Nat Rev Clin Oncol 19, 37–50 (2022).

47. Zhao, J. et al. Enhancing T cell anti-tumor efficacy with a PD1-TIGIT chimeric immune-checkpoint switch receptor. OncoImmunology 12, 2265703 (2023).

48. Robbins, P. F. et al. Cutting Edge: Persistence of Transferred Lymphocyte Clonotypes Correlates with Cancer Regression in Patients Receiving Cell Transfer Therapy. The Journal of Immunology 173, 7125–7130 (2004).

49. Wittibschlager, V. et al. CAR T-Cell Persistence Correlates with Improved Outcome in Patients with B-Cell Lymphoma. IJMS 24, 5688 (2023).

50. Chapuis, A. G., et al. Transferred melanoma-specific CD8+ T cells persist, mediate tumor regression, and acquire central memory phenotype. Proc. Natl. Acad. Sci. U.S.A. 109, 4592–4597 (2012).

51. Chapuis, A. G. et al. T-Cell Therapy Using Interleukin-21–Primed Cytotoxic T-Cell Lymphocytes Combined With Cytotoxic T-Cell Lymphocyte Antigen-4 Blockade Results in Long-Term Cell Persistence and Durable Tumor Regression. JCO 34, 3787–3795 (2016).

52. Kawalekar, O. U. et al. Distinct Signaling of Coreceptors Regulates Specific Metabolism Pathways and Impacts Memory Development in CAR T Cells. Immunity 44, 712 (2016).

53. Guedan, S., et al. Enhancing CAR T cell persistence through ICOS and 4-1BB costimulation. JCI Insight 3, e96976 (2018).

54. Carpenito, C., et al. Control of large, established tumor xenografts with genetically retargeted human T cells containing CD28 and CD137 domains. Proc. Natl. Acad. Sci. U.S.A. 106, 3360–3365 (2009).

55. Milone, M. C. et al. Chimeric Receptors Containing CD137 Signal Transduction Domains Mediate Enhanced Survival of T Cells and Increased Antileukemic Efficacy In Vivo. Molecular Therapy 17, 1453–1464 (2009).

56. Guedan, S., et al. Enhancing CAR T cell persistence through ICOS and 4-1BB costimulation. JCI Insight 3, e96976 (2018).

57. Lin, S. et al. Chimeric CTLA4-CD28-CD3z T Cells Potentiate Antitumor Activity Against CD80/CD86– Positive B Cell Malignancies. Front. Immunol. 12, 642528 (2021).

58. Guo, J.-X. et al. Bioactivity and safety of chimeric switch receptor T cells in glioblastoma patients. Front Biosci 24, 1158–1166 (2019).

59. Chimeric Switch Receptor Modified T Cells for Patients With PD-L1+ Recurrent or Metastatic Malignant Tumors. clinicaltrials.gov/study/NCT02930967 doi:NCT02930967.

60. Liu, H. et al. CD19-specific CAR T Cells that Express a PD-1/CD28 Chimeric Switch-Receptor are Effective in Patients with PD-L1–positive B-Cell Lymphoma. Clinical Cancer Research 27, 473–484 (2021).

61. Voss, R.-H., Kuball, J. & Theobald, M. Designing TCR for Cancer Immunotherapy. in Adoptive Immunotherapy vol. 109 229–256 (Humana Press, New Jersey, 2004).

62. Langlet, C., Neil, G. A. & Sherman, L. A. The mechanism of anti-Lyt-2 inhibition of antibody-directed lysis by cytotoxic T lymphocytes. J Immunol 139, 3590–3596 (1987).

