## Supplemental Material for "A signaling-optimized TIGIT chimeric switch receptor promotes metabolic fitness and antitumor efficacy of TCR/CAR T cells"

### Supplemental Methods

**Table 1: Primers**

| Usage | Target | Forward sequence | Reverse sequence |
|---|---|---|---|
| Sequencing | TIGIT/28 | TTACACAGTCCTGCTGACCACC | |
| Sequencing | TIGIT/28F | TTACACAGTCCTGCTGACCACC | |
| Site directed mutagenesis | TIGIT/28F | CACAGCGACTACATGTTCATGA CCCCTAGACG | CTGTGCAGCAGTCTGCTC |

**Table 2: Antibodies**

| Antigen | Fluorochrome | Isotype | Clone | Provider |
|---|---|---|---|---|
| **B7-H6** | PE | Mouse IgG1 | 875001 | R&D Minneapolis |
| **CD3** | APC | Mouse IgG2a, k | HIT3a | BD Bioscience |
| **CD4** | FITC | Mouse IgG1, κ | RPA-T4 | BD Bioscience |
| **CD4** | APC | Mouse IgG1, κ | RPA-T4 | BD Bioscience |
| **CD8** | FITC | Mouse IgG1, κ | RPA-T8 | BD Bioscience |
| **CD8** | PE-Cy7 | Mouse IgG1, k | RPA-T8 | Biolegend |
| **CD8** | APC | Mouse IgG1a, κ | RPA-T8 | BD Bioscience |
| **CD25** | PE | Mouse IgG1, k | BC96 | Biolegend |
| **CD27** | PE | Mouse IgG1, k | M-T271 | Biolegend |
| **CD28** | FITC | Mouse C3H x BALB/c IgG1, k | CD28.2 | BD Bioscience |
| **CD28** | PE | Mouse C3H x BALB/c IgG1, k | CD28.2 | BD Bioscience |
| **CD45RA** | PE | Mouse IgG2b, κ | HI100 | BD Bioscience |
| **CD45RA** | V450 | Mouse IgG1, κ | 5H9 | BD Bioscience |
| **CD57** | APC | Mouse IgM, κ | NK-1 | BD Bioscience |
| **CD62L** | PE-Cy5 | Mouse IgG1, κ | DREG56 | Beckman Coulter |
| **CD107A** | PE-Cy5 | Mouse BALB/c IgG1, k | H4A3 | BD Bioscience |
| **CD155** | PE | Mouse BALB/c IgG1, k | SKII.4 | BD Bioscience |
| **CD160** | PE | Mouse BALB/c IgM | BY55 | BD Bioscience |
| **CD226** | FITC | recombinant human IgG1 | REA1040 | Miltenyl |
| **LAG-3** | BV421 | Mouse IgG1, κ | T47-530 | BD Bioscience |
| **TIM-3** | BV421 | Mouse IgG1, κ | 7D3 | BD Bioscience |
| **TIGIT** | PE | recombinant human IgG1 | REA1004 | Miltenyi Biotech |
| **TIGIT** | APC | recombinant human IgG1 | REA1004 | Miltenyi Biotech |
| **PD-1** | FITC | Mouse IgG1, κ | MIH4 | BD Bioscience |
| **PD-1** | APC | Mouse IgG1, κ | EH12.2.H7 | BD Bioscience |
| **PD-L1** | APC | Mouse IgG1, κ | MIH1 | BD Bioscience |
| **TCR Vβ3** | PE | recombinant human IgG1 | REA646 | Miltenyi Biotech |

**Cell line, peptides and cytotoxicity assay**

The HLA_A2.1$^{+}$ p53 null osteosarcoma cell line Saos2 was maintained in RPMI (GIBCO) supplemented with 10% FCS, 1% glutamine, and 1% penicillin-streptomycin. HLA.A2.1-restricted p53(264–272)[1, 2] peptide was synthesized by Biosyntan (Berlin, Germany) and resuspended in DMSO at the indicated concentration. Saos2 pulsed with the cognate peptide to determine EC50 values was performed in duplicate wells as reported previously[3]. Standard $^{51}$Cr-release assays were performed at the indicated effector-to-target (E:T) ratio in duplicate wells and the half-maximal stimulatory concentration (EC50) of the different experimental groups were determined with Prism version 10.0 (GraphPad Software) as described earlier[3].

#### Supplementary references

1.Theobald M., Biggs J., Dittmer D., Levine A.J., Sherman L.A. Targeting p53 as a general tumor antigen. Proc. Natl. Acad. Sci. USA. 1995;92:11993–11997. doi: 10.1073/pnas.92.26.11993.

2.Theobald M., Ruppert T., Kuckelkorn U., Hernandez J., Häussler A., Ferreira E.A., Liewer U., Biggs J., Levine A.J., Huber C. The sequence alteration associated with a mutational hotspot in p53 protects cells from lysis by cytotoxic T lymphocytes specific for a flanking peptide epitope. J. Exp. Med. 1998;188:1017–1028. doi: 10.1084/jem.188.6.1017.

3. Echchannaoui H, Petschenka J, Ferreira EA, Hauptrock B, Lotz-Jenne C, Voss RH, Theobald M. A Potent Tumor-Reactive p53-Specific Single-Chain TCR without On- or Off-Target Autoimmunity In Vivo. Mol Ther. 2019 Jan 2;27(1):261-271. doi: 10.1016/j.ymthe.2018.11.006.

### Supplemental Figures

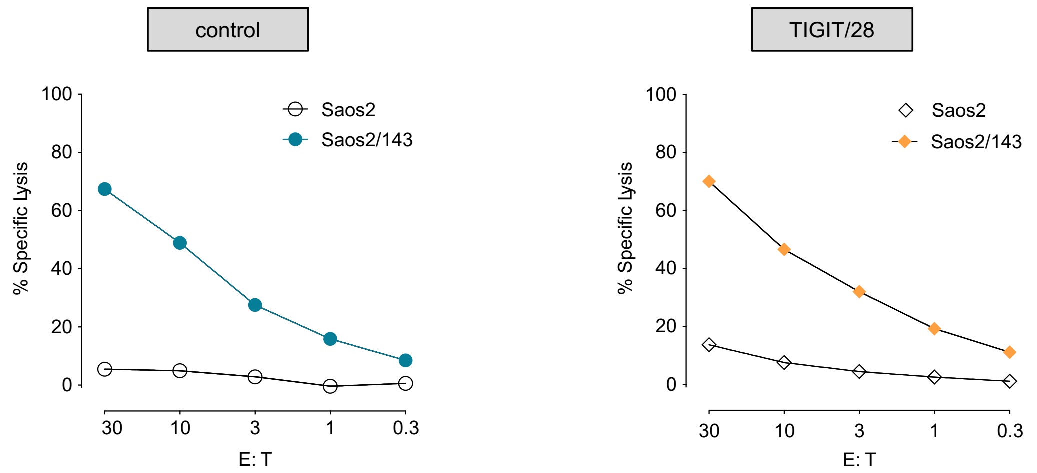


**Supplemental-Figure 1. Short-term cytolytic activity of TIGIT/28 T cells.** $^{51}$Cr-release assay was used to evaluate short-term antitumor responses of p53TCR (control) and p53TCR/CSR (TIGIT/28) modified T cells. Specific lysis of p53$^{+}$A2.1$^{+}$ target (Saos2/143) and p53$^{-}$A2.1$^{+}$ (Saos2) control tumor cells after 5 h co-culture with control or TIGIT/28 T cells is illustrated at the indicated effector to target ratio (CD8$^{+}$Vβ3$^{+}$:Tumor).

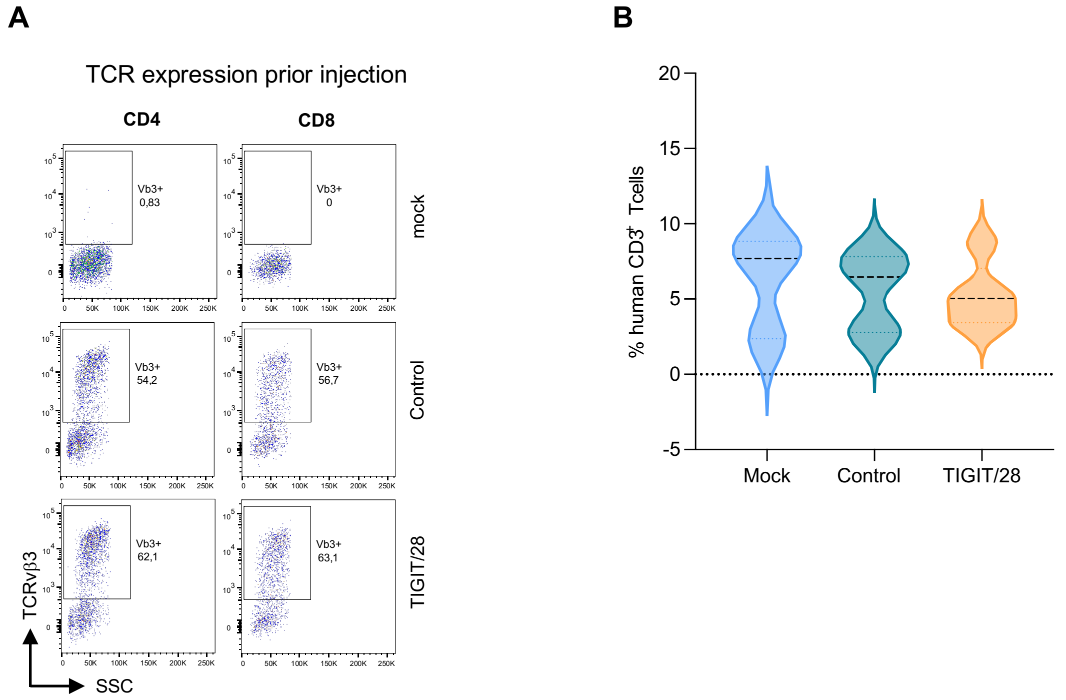


**Supplemental-Figure 2. Characterization of TIGIT/28 CSR T cells in xenograft tumor model.** Experimental design as described in Figure 2D. **A)** Flow cytometry pseudocolor plots demonstrating the frequency of CD4$^{+}$ and CD8$^{+}$ T cells and surface expression of Vβ3 before injection into NSG mice of mock, control and TIGIT/28 T cells. **B)** Frequency of human CD3$^{+}$ T cells in the peripheral blood of mice receiving mock, control or TIGIT/28 T cells, measured by flow cytometry at day 22 (day 14 after T cell injection) (n=3-5 animals).

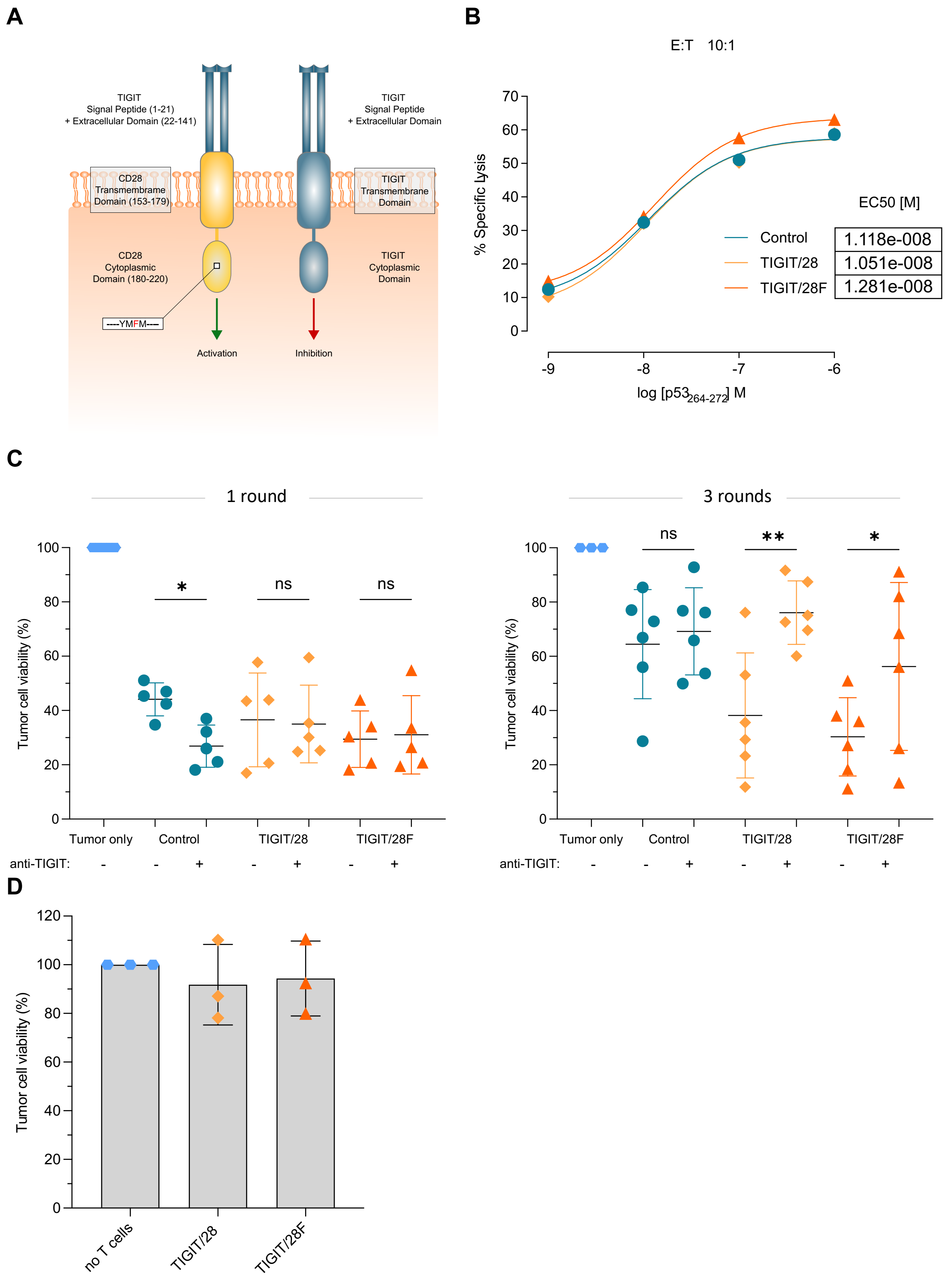


**Supplemental-Figure 3. Functional characterization of the optimized TIGIT/28F CSR in vitro. A)** Graphical illustration of the optimized CSR design showing the amino acid change from asparagine to phenylalanine at position 182 to turn the YMNM motif of CD28 to the YMFM motif. **B)** Half maximal effective peptide concentration (EC50) of the applied p53TCR in CSR TIGIT/28, TIGIT/28F or control T cells as determined by $^{51}$Cr-release assay. EC50 was measured based on the specific lysis of the p53$^{-}$ A2.1$^{+}$ (Saos2) pulsed with titrated amount of p53$_{(264-272)}$ peptide at the indicated E:T ratio. The percentages of lysis are calculated as means from technical duplicates and shown for a representative chromium release assay. **C)** Tumor cell viability (Saos2/143) after one round (24 h) and three rounds (3 x 24 h) long-term tumor colony-forming assay of TIGIT/28, TIGIT/28F or control T cells with and without a monoclonal anti-TIGIT blocking antibody (1 round: Control $p$ = 0.02, TIGIT/28 $p$ = 0.996, TIGIT/28F $p$ = 0.9946, 3 rounds: Control $p$ = 0.7088, TIGIT/28 $p$ = 0.0074, TIGIT/28F $p$ = 0.0448, n=5 and n=6 biological replicates). Statistical significance was determined by one-way ANOVA and Šidák's multiple comparisons test. *$P < 0.05$; **$P < 0.01$; ***$P < 0.001$; ns, not significant. **D)** Viability of $p53^+A2.1^+$ (Saos2/143) tumor cells determined by long-term tumor colony forming assay after co-culture with T cells expressing TIGIT/28 or TIGIT/28F CSR without p53TCR. Experimental condition containing tumor cells only (without co-culture with T cells) is used as negative control. Lysis values were normalized to control group (n=3 biological replicates).

A

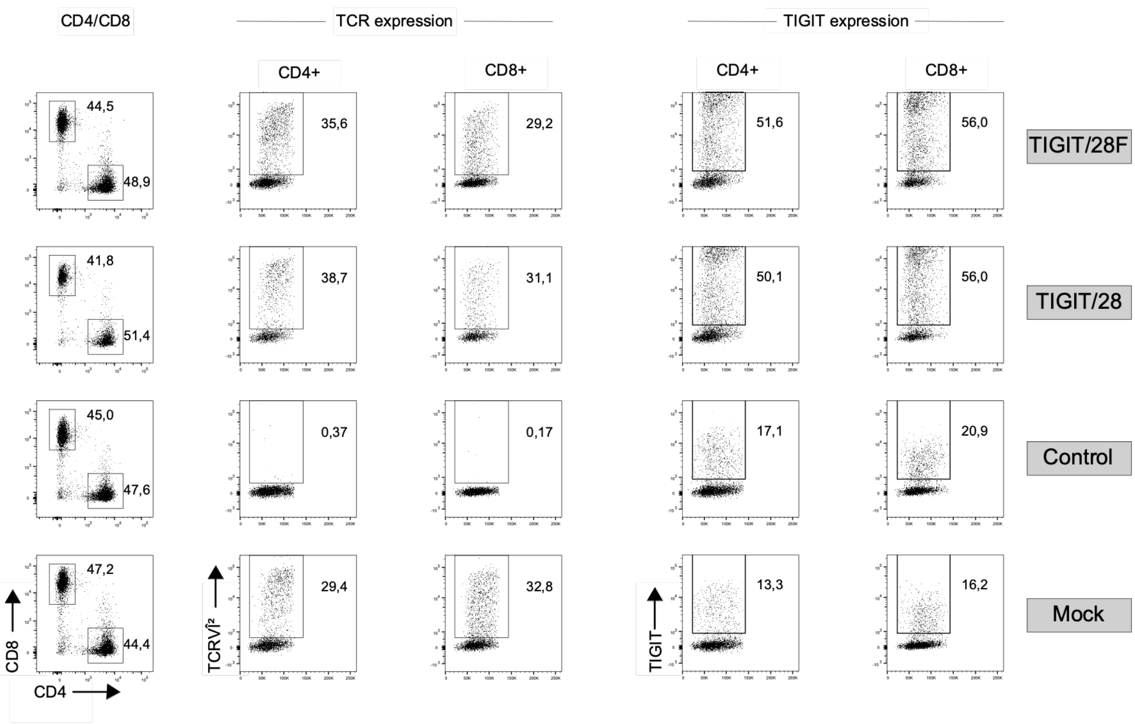

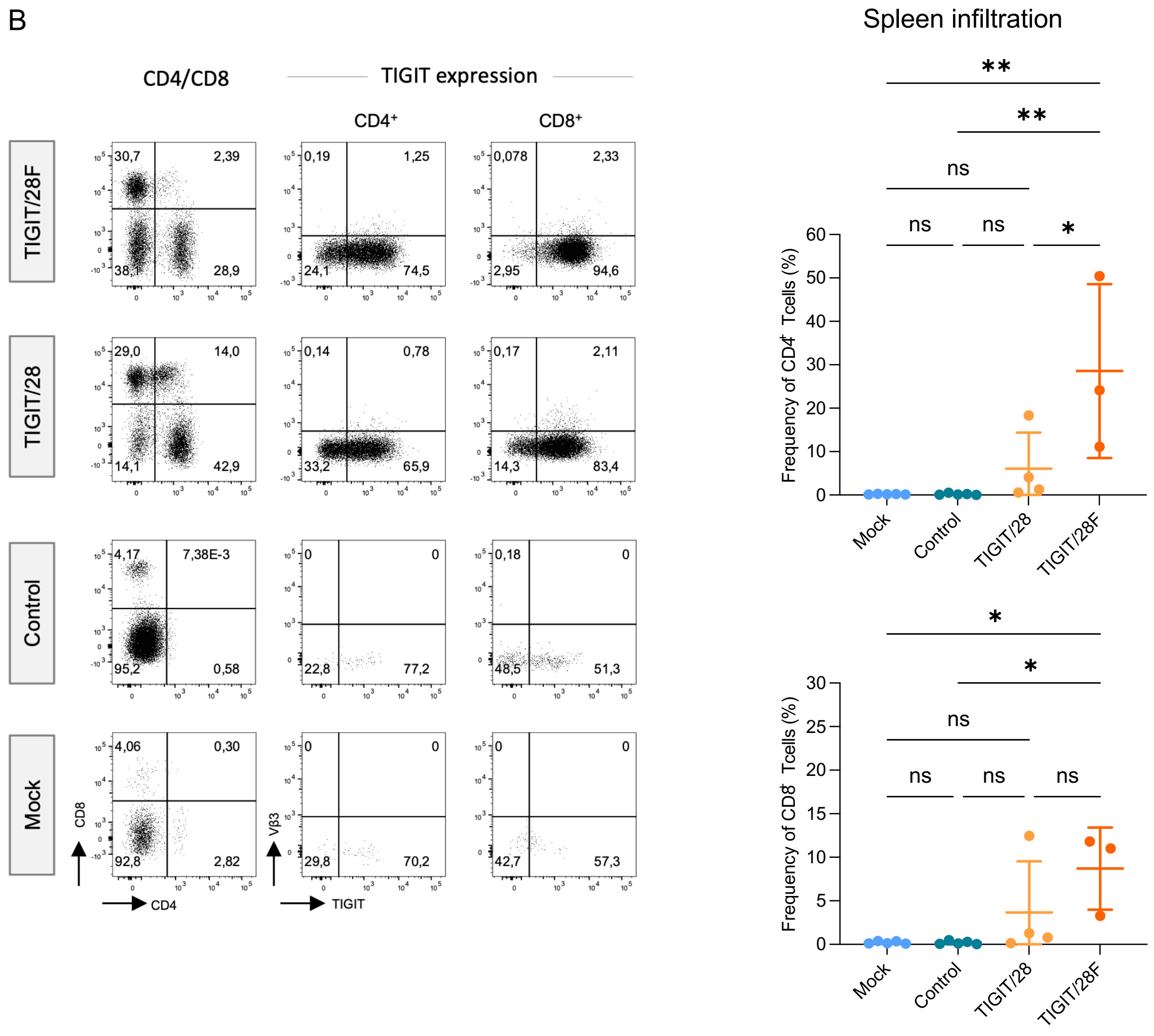


**Supplemental-Figure 4. Evaluation of the anti-tumor function of the optimized TIGIT/28F CSR in vivo.** Experimental design of the xenograft model to evaluate anti-tumor responses *in vivo* is already described in the main Figure 4A. **A)** Flow cytometry plots depicting the frequency of CD4⁺ and CD8⁺ T cells and their respective surface expression of TCRVβ3 and TIGIT prior injection into NSG mice. Data are shown for mock, control, TIGIT/28 and TIGIT/28F CSR T cells. **B)** Murine spleen cells were isolated (from sacrificed animals reaching human endpoints) and the frequency of infiltrating human CD4⁺ and CD8⁺ T cells expressing endogenous TIGIT or CSR in mice treated with mock, control or TIGIT/28 and TIGIT/28F CSR T cells was analysed by flow cytometry. Representative flow cytometry plots (left) and quantification from all mice (right) are shown. Statistical significance was determined by ordinary one-way ANOVA and Tukey's multiple comparisons test. **P* < 0.05; ***P* < 0.01; ****P* < 0.001; ns, not significant (n=3-5 animals/group).

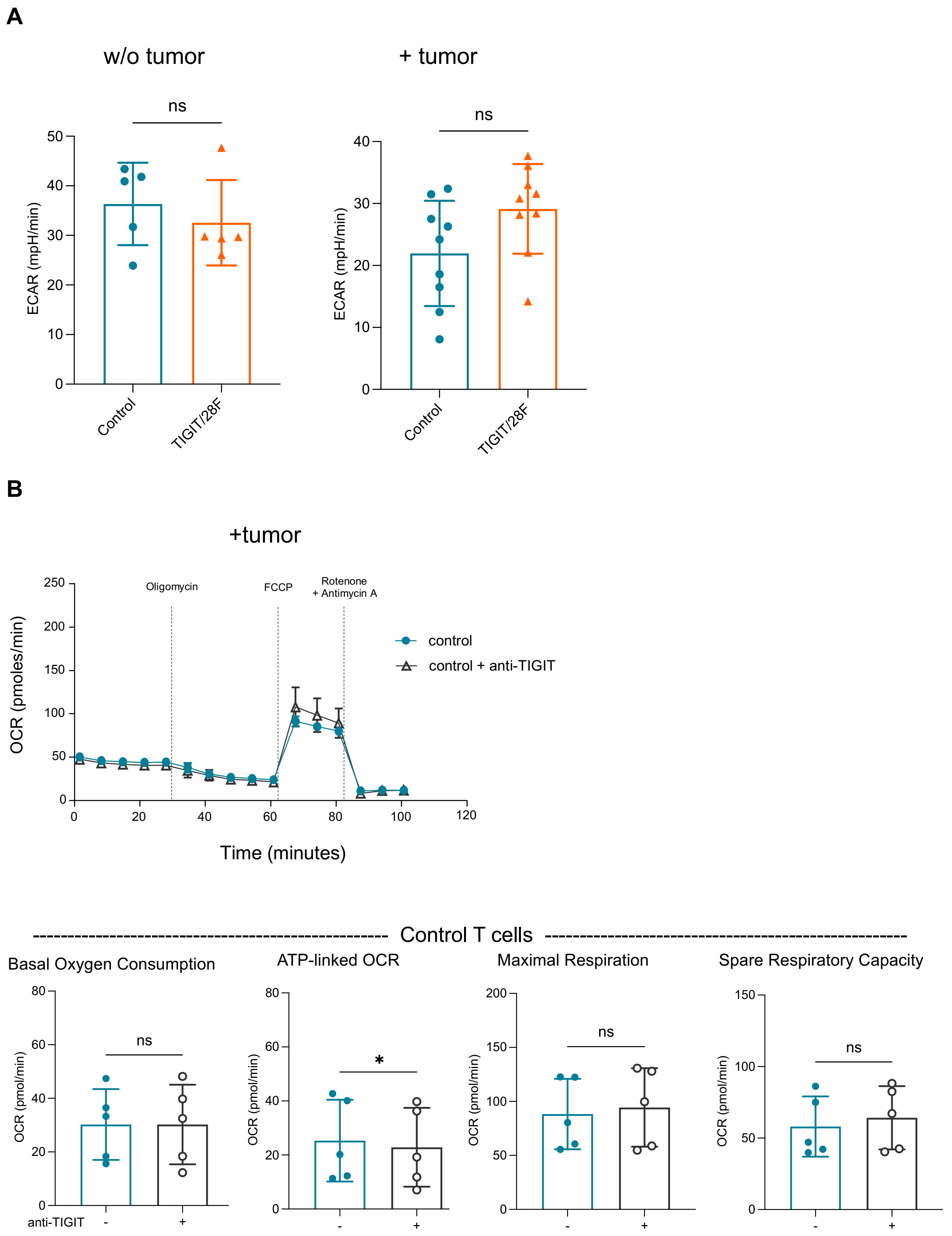
A
w/o tumor
+ tumor
ns
ECAR (mpH/min)
Control
TIGIT/28F
B
+tumor
Oligomycin
FCCP
Rotenone
+ Antimycin A
control
control + anti-TIGIT
OCR (pmoles/min)
Time (minutes)
Control T cells
Basal Oxygen Consumption
ATP-linked OCR
Maximal Respiration
Spare Respiratory Capacity
OCR (pmol/min)
ns
*
anti-TIGIT
-
+

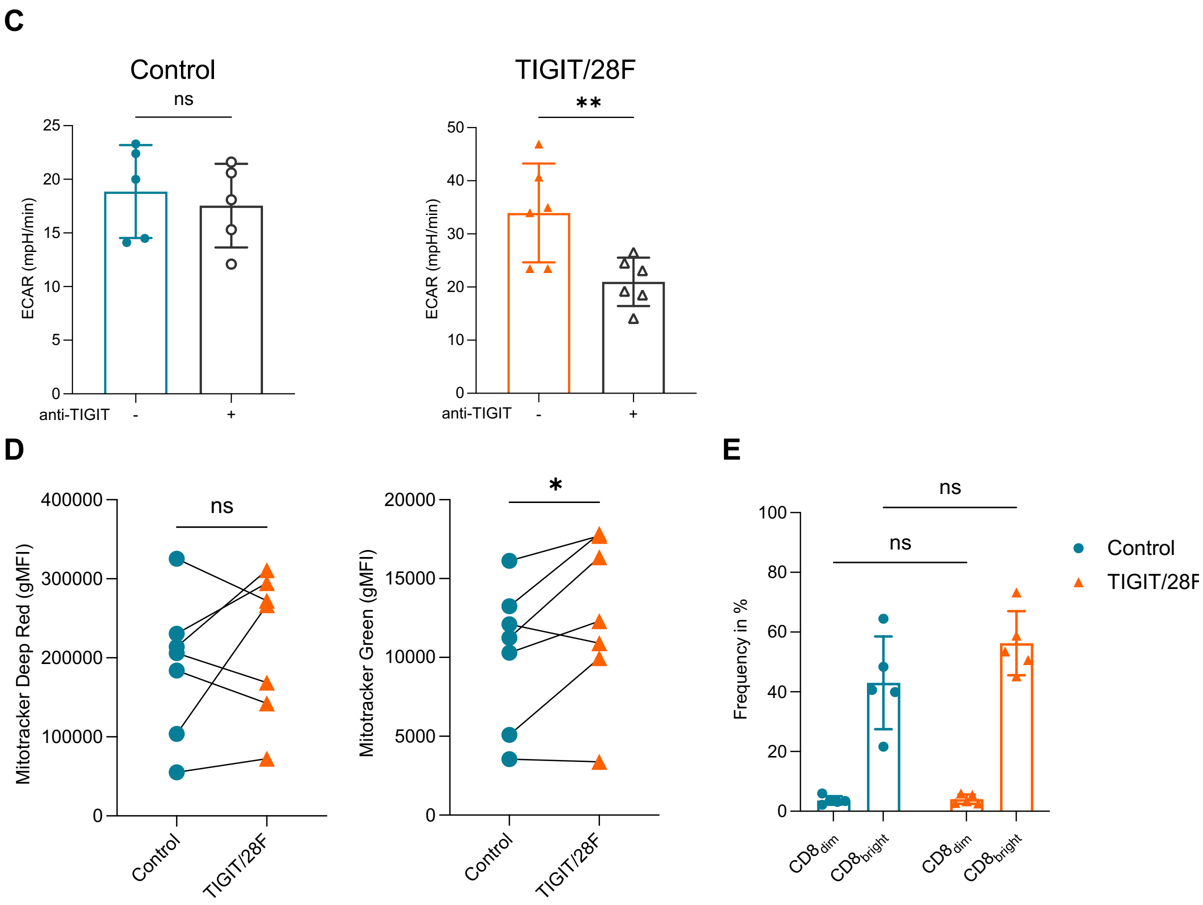


**Supplemental-Figure 5. Analysis of metabolic activity in TIGIT/28F CSR T cells upon co-culture with target tumor cells. A)** Cumulative data of basal extracellular acidification rate (ECAR) of control and TIGIT/28F T cells from *in vitro* culture after 48h co-culture with target tumor cells Saos2/143 (Control vs. TIGIT/28F $p$ = 0.074 or without tumor cells as control $p$ = 0.4224), measured by Seahorse XFp extracellular flux assay. **B)** Example of Seahorse XFp extracellular flux 'Mito stress test' (top) and cumulative data (bottom) of basal ($p$ = 0.9999) and ATP-linked oxygen consumption rate ($p$ = 0.0287), maximal ($p$ = 0.185) and spare respiratory capacity ($p$ = 0.1826) of control T cells after co-culture with target tumor cells (48h) +/- anti-TIGIT blocking mAb. **C)** Cumulative data of basal ECAR from control (left, $p$ = 0.0682) and TIGIT/28F (right, $p$ = 0.0029) T cells after co-culture with target tumor cells (48h) +/- anti-TIGIT blocking mAb. **D)** Mitochondrial membrane potential (left, $p$ = 0.3714) and mass (right, $p$ = 0.0463) indicated by flow cytometric gMFI of MitoTracker Deep red and MitoTracker Green staining of total $CD8^+$ T cells. **E)** Frequency of $CD8^{dim}$ ($p$ = 0.9971) and $CD8^{bright}$ ($p$ = 0.0819) cells from flow cytometric measurements of control and TIGIT/28F T cells. Statistical significance in A-D was determined by two-tailed Student's $t$-test and in E by two-way ANOVA followed by Šidák's multiple comparisons test. *$P < 0.05$; **$P < 0.01$; ***$P < 0.001$; ns, not significant. n=5-9 biological replicates.

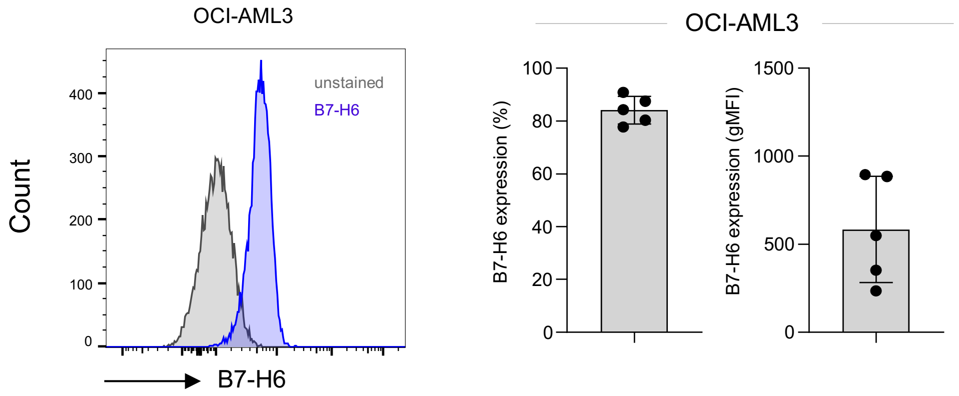


**Supplemental-Figure 6. Analysis of surface expression of the CAR ligand (B7-H6) in OCI-AML3 tumor cell line.** Representative histogram (left) and cumulative data (right) showing surface expression of the NKp30 CAR target B7-H6 (in percent and gMFI) in OCI-AML3 tumor cells, as determined by flow cytometry (n=5 repeated measurements).
